# Intragastric administration of low-dose rotenone post-colitis exacerbates damage to the nigrostriatal dopaminergic system in Parkinson’s disease: The pace accelerates even more

**DOI:** 10.1101/2022.12.22.521569

**Authors:** Nishant Sharma, Monika Sharma, Disha Thakkar, Hemant Kumar, Sona Smetanova, Lucie Buresova, Petr Andrla, Amit Khairnar

**Affiliations:** Department of Pharmacology and Toxicology, National Institute of Pharmaceutical Education and Research (NIPER), Ahmedabad, Gujarat, India; Department of Pharmaceutical Analysis, National Institute of Pharmaceutical Education and Research (NIPER), Ahmedabad, Gujarat, India; RECETOX, Faculty of Science, Masaryk University, Brno, Czech Republic; International Clinical Research Centre, St. Anne’s University Hospital, Brno, Czech Republic, ICRC, FNUSA, Brno, Czech Republic

**Keywords:** Parkinson’s disease, Gut inflammation, Inflammatory bowel disease, Alpha synuclein progression, Gut brain axis

## Abstract

**Background:** The contribution of gastrointestinal (GI) inflammation and local exposure to neurotoxins in the gut offers the most in-depth explanation of Parkinson’s disease (PD) etiopathogenesis through abnormal accumulation and spreading of alpha-synuclein (α-syn) aggregates from the gut to the brain.

**Objectives:** This study was designed to investigate whether dextran sodium sulfate (DSS)-mediated colitis may have lasting effects on dopaminergic pathways in the brain and whether or not colitis exacerbated susceptibility to later exposure to the neurotoxin rotenone.

**Methods:** To induce chronic colitis, 10 months old C57BL/6 mice were pre-exposed to 3 cycles of 7 days of 1% (w/v) DSS administration in drinking water followed by 14 days of regular drinking water. After colitis-induction, animals received a low dose of intragastric rotenone for the next 8 weeks, followed by testing for Parkinsonian behavior and GI phenotypes of inflammation. At the end of the 8^th^ week after colitis, colon, brain stem, and midbrain tissue were isolated and analyzed for α-syn, inflammatory markers, and dopaminergic neuronal loss. Gut microbial composition was assessed by 16S rRNA sequencing analysis.

**Results:** We found that local rotenone exposure for 8 weeks did not affect colitis severity and colonic tight junction(TJ) protein expression (ZO-1, Occludin, and Claudin-1). On the other hand, we found that while eight weeks of chronic rotenone administration led to an increase in inflammatory markers, the presence of pre-existing colitis resulted in a considerable change in gut microbiota composition and a decrease in TJ’s protein expression. In addition, the administration of rotenone in mice post-colitis caused gastrointestinal function impairment and poor behavioral performances. Itworsened rotenone-induced α-syn pathology in the colon, which extended upward and resulted in severe dopaminergic neuron loss and significant astroglia activation in the dorsal motor nucleus of the vagus (DMV), locus coeruleus, substantia nigra as well as in striatum. Interestingly, in the case of rotenone alone, we found that α-syn induced ChAT^+^ neuronal death is restricted to the DMV. These findings indicate that long-term rotenone exposure in conjunction with early inflammatory intestinal milieu exacerbates the progression of α-syn pathology and aggravates neurodegeneration in the intragastric mouse PD model.

**Conclusions:** This work provides detailed insight into the involvement of GI inflammation triggered after a neurotoxic insult in the colon and explores their potential to impact central dopaminergic degeneration in PD. This way, we can identify potential therapeutic targets that stop the enteric inflammatory processes involved in progressing PD.

**Graphical Abstract:** 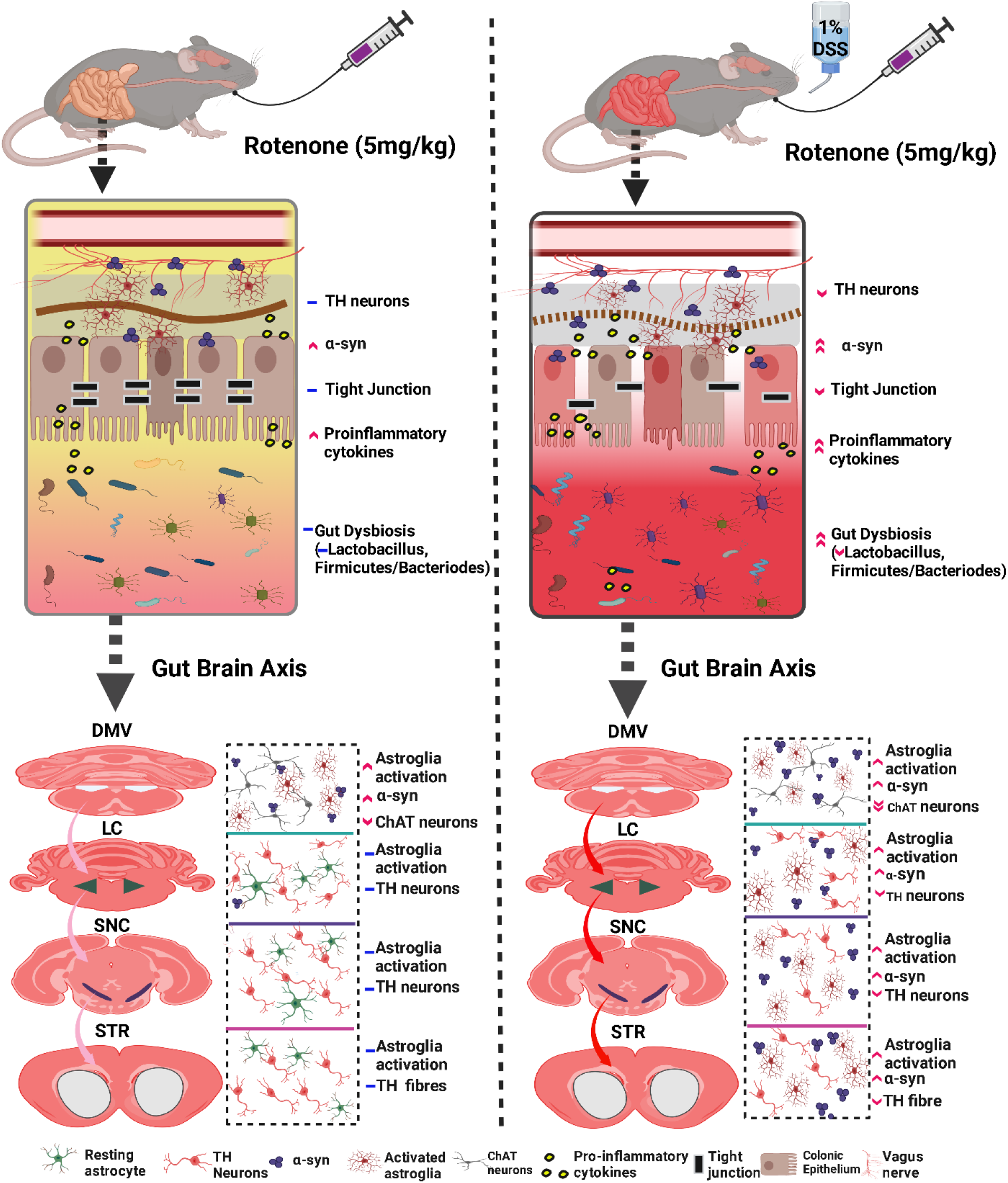

**Highlights:**

- Low-dose intragastric rotenone post-colitis aggravates gastrointestinal dysfunction and accelerates the onset of motor impairment.
- Low-dose intragastric rotenone did not alter colitis clinical and histological aspects.
- Low-dose intragastric rotenone post-colitis exacerbated the progression of α-syn pathology from the GI tract to the brain, leading in central dopaminergic neuronal degeneration.

## 1. Introduction

Parkinson’s disease (PD) is not just reflective of synucleinopathies that affect the brain but also the gastrointestinal (GI) tract^1^. The pathological features of PD include the progressive loss of dopaminergic (DAergic) neurons within substantia nigra pars compacta (SNpc), as well as the appearance of intraneuronal inclusions known as Lewy bodies or Lewy neurites, which are primarily composed of self-aggregatory protein alpha-synuclein (α-syn)^2^. Current epidemiological research suggests that PD patients have a variety of prodromal GI dysfunctions, including delayed gastric emptying, infrequent bowel movements, and chronic constipation, decades before they acquire centrally driven motor deficits of PD^3^. Correspondingly, GI tissue of PD patients has shown changes in functional and structural activities during the early stage of the disease, including the presence of α-syn containing inclusions, impaired mucosal barrier function, and altered bacterial composition^4,5.^ The presentation of GI pathology is linked with the Braak hypothesis, which states that ingestion/inhalation of unknown environmental triggers causes the development of α-syn pathology first in the enteric nervous system (ENS), followed by its propagation to lower brain stem regions, such as the dorsal motor nucleus of the vagus nerve (DMV), locus coeruleus (LC) and subsequently to the SNpc, where it induces death of DAergic neurons^6^. In support of this, several preclinical studies have recently demonstrated transneuronal propagation of injected recombinant α-syn into the gut wall to the brain *via* the vagal nerve^7–9^. Clinical studies have also revealed that complete truncal Vagotomy is now linked to a lower risk of PD^10^. These reports indicate that enteric α-syn pathology can spread from the GI tract to the brain. Still, only a few studies have been conducted to explain the molecular mechanism associated with the development of enteric α-syn pathology and its long-distance upward transmission to the brain.

As a result, several speculations have been about the involvement of early intestinal inflammation in PD pathogenesis. Indeed, recent population-based cohort studies have shown that people with inflammatory bowel disease (IBD) are more likely to get PD later in life, and blocking the tumor necrosis factor-α (TNF-α) pathway lowers this risk^11,12.^ IBD is an umbrella term for Crohn’s disease and ulcerative colitis, both of which are characterized by the chronic inflammation of the GI tract^13^. Indeed, α-syn levels were found to be elevated throughout the enteric neurons of children with intestinal inflammation and adults with Crohn’s disease ^14,15.^ Consequently, individuals with IBD are exposed to many micro-environmental factors throughout their lifetime. Environmental factors, including pesticides, have been proven to impair the host immune response in the GI tract, limit beneficial bacterial cells and alter mucosal barrier function^16^. These overlapping molecular and biological mechanisms may help to uncover the common pathogenic pathways for both diseases. A more likely scenario, therefore, involves repeated exposures to low doses of toxins, whose pathogenicity may be enhanced by early intestinal inflammation.

Moreover, these two seemingly disparate diseases also share common genes mutations, such as leucine-rich repeat kinase 2 (LRRK2), which is now associated with causing PD-associated neuroinflammation and neuropathology in response to dextran sodium sulfate (DSS) induced colonic inflammation in LRRK2 G2019S transgenic mice ^17^. DSS is the most common colitogen that induces colitis in rodents by disrupting the intestinal epithelial barrier and stimulating colonic inflammation due to an overall shift in quantity and diversity of microbiota constituents^18^. Furthermore, activation of enteric glia and greater expression of genes encoding pro-inflammatory mediators, including TNF-α, interleukin -6 (IL-6), interleukin-1β (IL-1β), and C-C chemokine ligand 2 (CCL2) has been observed in PD patients’ colon biopsies as compared to age-matched normal subjects^19^. However, it is currently unknown whether early inflammation or inflammation along with exposure to environmental toxins in the GI tract may promote the formation of α-syn accumulation and subsequent development of PD in genetically susceptible individuals.

The possible contribution of suspected environmental triggers in the association between early GI inflammation and the progression of PD has received little consideration. From this perspective, it is reasonable to presume that ingestion of a common environmental toxin may cause PD indirectly by promoting GI inflammation, hence encouraging α-syn misfolding accumulation in the ENS. However, there is a shortage of animal models that mimic the complex and progressive neuronal characteristics of PD, particularly about long-term environmental toxin exposure and early GI inflammation, that are needed to investigate this slow and progressive PD-related neurodegeneration and to develop specific treatment strategies.

The objective of this study is to examine two pertinent topics to advance our understanding of the role of intestinal inflammation in PD. To determine if the simultaneous presence of early GI (colonic) inflammation and compromised intestinal barrier integrity contributes to the course of disease in an intragastric rotenone mouse PD model. Second, we examined the association between chronic GI inflammation and the induction of PD pathophysiology in the presence/absence of environmental contaminants. This research has the potential to inform and direct population-level efforts to minimize exposure to environmental toxins that have been linked to the development of PD, as well as to help individual IBD patients reduce their own risk.

## 2. Material and methods

### 2.1. Materials

Dextran sodium sulfate (molecular weight: 40,000kDa), Rotenone, Acrylamide, Sodium dodecyl sulfate (SDS), Tween-20, Formaldehyde, 3,3′-diaminobenzidine (DAB), glucose oxidase, D-glucose were obtained from Sigma Aldrich (MA, USA). TNF-α and IL-6 enzyme immunoassay kits were procured from Abcam (MA, USA). The protein ladder and PVDF membrane were obtained from Bio-rad (CA, USA). HRP conjugated Enhanced Chemiluminescent Substrate Reagent kit was procured from Invitrogen. All primary and secondary antibodies were procured from Abcam, Sigma, and Merck.. The Vectastain ABC kit was procured from Vector Laboratories (CA, USA) for immunohistochemical staining.

### 2.2. Animals and experimental protocol

Male C57BL/6J mice (n=40, 10 months old, 28-33 g) were procured from the Zydus Research center Ahmedabad. We chose to use aged male mice because the incidence of PD is biased toward males and the aging factor^20^. All mice were randomly allocated to five per cage and housed in a climate-controlled room with a temperature of 24 ± 2 °C and an indoor relative humidity of 50 ± 10% (12 hours light/dark cycle). Mice had ad libitum access to food and water. All animals experiments were approved by the Institutional Animal Ethical Committee (IAEC/2018/031) of the National Institute of Pharmaceutical Education and Research (NIPER), Ahmedabad, India, (Regd. No.1945/GO/Re/s/17CPCSEA) and were conducted according to Committee for the Purpose of Control and Supervision of Experiments on Animals (CPCSEA) guidelines, India.

This experimental design consisted of seventeen weeks (Figure 1A). Mice underwent acclimatization and baseline testing for behavioral tests for two weeks before starting the experiments. Animals were randomly allocated into four groups. Group I: Control mice received 0.5% carboxymethyl cellulose (CMC) and sterile H2O. Group II: DSS alone (DSS) mice received 1% DSS for 3 cycles. After induction of colitis, these mice were administered 0.5% CMC from the 9^th^ week till the 17^th^ week of the study. Group III: Rotenone alone (Rot) mice in this group first received sterile water from weeks 0 to 9^th^. These mice received a low dose of intragastric rotenone (5mg/kg, for 5 days/week) starting from the 9^th^ week till the 17^th^ week of the study. Group IV: DSS+Rotenone (DSS+Rot) mice received 1% DSS in drinking water for 3 cycles to induce chronic colitis. These mice were later administered a low dose of intragastric rotenone (5mg/kg/day, for 5 days/week) for the next 8 consecutive weeks (9^th^ to 17^th^week). In the second and fourth groups, chronic colitis in mice was induced by giving 1% DSS in drinking water for 3 consecutive cycles. Each cycle comprised of administration of 1% DSS for 7 days, followed by recovery with sterile water for the next 2 consecutive weeks, and this was defined as one cycle of DSS^21,22.^ Control and Rot alone group animals received clean water throughout. Rotenone is freely soluble in chloroform. A stock solution of 50 mg of rotenone in 1 ml of chloroform was prepared and stored at −20°C for not more than two weeks. Testing for Parkinsonian behavior and GI phenotypes of inflammation were accessed at different time points. Mice were weighed on a weekly basis. During the study, Serum was collected at various time points (9th, 13th, and 17th week). To assess the gut microbiota composition, 16S rRNA sequencing was performed on fecal samples collected in the 17th week (Figure 1A).

**Figure 1:**
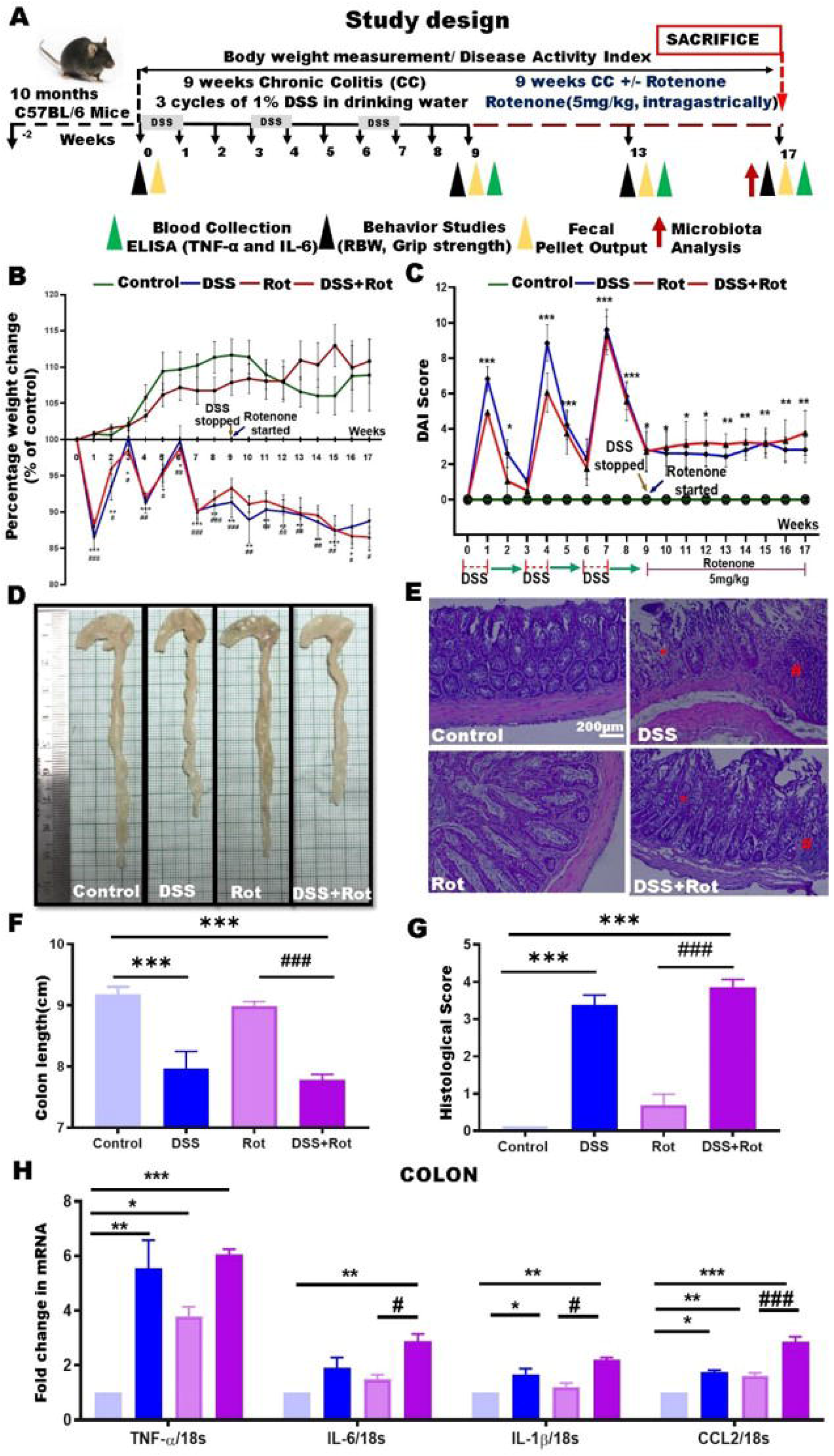
A) Experimental procedure followed by combining two experimental models: chronic intestinal inflammation caused by 1% DSS in drinking water and an intragastric rotenone mouse model of PD B) Body weight, as a percent of starting weight, assessed weekly (n=8-12). C) Disease activity index reflecting weight loss, loose stool consistency, and amount of blood present in stool and rectum, assessed throughout the recurring DSS-induced colitis and rotenone administration time course every 7 days(n=8-12). D, F) Colon length 17^th^ week (n=5). E, G) Representative H&E-stained middle colonic sections and histological scores were assessed at the end of the 17^th^ week (magnification, x100) (n=5). In DSS and DSS+Rot, an asterisk (*) indicates an area of goblet cell depletion and distortion of crypt architecture. (#) indicate crypt abscesses with neutrophils and damaged epithelial cells in cross-section (×10, scale bar 200 μm). H) Relative mRNA expression of inflammatory cytokines TNF-α, IL-6, IL-1β, and CCl2 in the colon was determined by real-time PCR (n=4-5). Data were expressed as Mean±SEM. *p< 0.05 **p< 0.01 ***p< 0.001 vs. control. ^#p^< 0.05 ^##^p< 0.01, ^###^p< 0.001 vs. rotenone. Statistical analysis was performed using one and two-way analysis of variance (ANOVA) followed by Tukey’s multiple comparison test.

### 2.3. Post-colitis rotenone measurement in the Serum, brain stem, and brain: A pilot study

Using liquid chromatography-mass spectroscopy (LC-MS), we determined the dose of rotenone post-colitis that was not detectable in Serum, brain stem, and in the brain of mice. In brief, eight-week-old C57BL/6 male mice (n=4) were treated with 3 cycles of 1% DSS, followed by administration of different doses of rotenone (1.25, 2.5, 5, 7.5 mg/kg) for 1 week. On the 7^th^ day, 200-250μl of blood was collected from the retro-orbital plexus at 1.5 and 3 h after rotenone administration and pooled in the same tube. Blood samples were then allowed to clot by leaving them undisturbed at room temperature for 30 minutes, and serum samples were isolated by centrifugation at 3000 rpm for 10 minutes. Mice were then sacrificed, and the brain and the brain stem were isolated from each mouse and stored at −80°C for further analysis. Rotenone was extracted from Serum, brain, and brain stem homogenates using methanol by protein precipitation method. Samples were analyzed using LC-QTOF-MS instrument, central instrumental facility, NIPER-Ahmedabad as previously described^23^. The LC-MS methods were linear over a concentration range from 10 to 1000 ng/ml. The limit of detection was found to be 10 ng/ml.

### 2.4. Behavioral tests

Two different behavioral experiments were conducted at four other time points (0, 9^th^, 13^th^, and 17^th^ week), including 1. Round beam walk (RBW) test for motor impairment: During this test, mice were trained on different-width beams. In the first attempt, a 25 mm-diameter beam was mounted 50 cm above the tabletop as the start platform and home cage. Each time mouse was trained to go home cage from the start platform. For the second trial, a medium-sized beam was used (15 mm in diameter). A video camera was put opposite the animal’s home cage to evaluate motor activity. A blind evaluator measured the time taken to cross the beam and the number of slips from the video. Detailed procedures for these tests were described previously^24^ 2. A grip strength test was used to assess muscle strength: During this procedure, the mice were placed on a planar square grid (15 cm**^2^**) with a wire mesh of (0.5 cm**^2^**). Once the mouse held the grid, the mouse was pulled back by holding the tail. The maximum force to keep the grid was recorded in grams. Three trials were performed for each mouse with a 30 minutes interval between tests ^20^.

### 2.5. Evaluation of Gut Pathology

#### 2.5.1. Disease Activity Index (DAI)

The DAI score was determined by combined scores of 1) Body weight loss, 2) Stool consistency, and 3) Fecal occult blood. Each score was determined based on loss in body weight, stool consistency, and blood in the stool as previously described (Table 1)^25^. Fecal occult blood testing of stool samples was performed using benzidine reagent^26^. Body weight loss was calculated as the percentage difference between the original body weight (day 0) and the body weight on any measurement day. At the end of the experiment, all mice were sacrificed, and the colon was separated from the vermiform appendix to the anus. The colon length was measured between the caecum and proximal rectum.

**Table 1:** Evaluation of Disease Activity Index.

| Stool Score | Weight loss | Stool consistency | Occult/gross bleeding |
| --- | --- | --- | --- |
| 0 | None or gain | Formed and hard | Absence |
| 1 | 1-5 | Formed but soft |  |
| 2 | 5-10 | Loose stool | Presence |
| 3 | 10-15 | Mild diarrhea |  |
| 4 | >15 | Gross diarrhea | Gross bleeding |

#### 2.5.2. Measurement of colon motility by one-hour stool collection

Fecal pellet output testing was performed on mice at four different time points (0, 9^th^, 13^th^, 17^th^ week). After fasting for 2 h, each mouse was placed in clean cages with no bedding, food, or water. After one hour in the new habitat, we collected the fecal pellets and counted the numbers. Subsequently, the wet weight was obtained by weighing the fresh pellets in preweighed tubesBy drying the pellets overnight at 65°C in an oven and reweighing the tubes to get dry weight. The water content percentage was determined based on the difference in the wet and dry weights of fecal pellets^27,28.^

### 2.6. Quantification of cytokines level (TNF-α, IL-6) in Serum

Blood samples were collected into 500μl EDTA tubes at different time points. Blood samples were then allowed to clot by leaving them undisturbed at room temperature for 30 minutes. The clot was removed by centrifuging at 3000 x g for 10 minutes at 4°C. Pale yellow supernatant was isolated from the tubes and used for TNF-α, IL-6 ELISA. The cytokines level was quantified using TNF-α (Abcam, 208348) and IL-6 (Abcam, 46100) according to manufacturer’s protocol.

### 2.7. Tissue collection and processing

At the end of the 17^th^ week of the experimental period, the first set of twenty mice (n=5 animals per group) sacrificed by decapitation using isoflurane. Brain and colon parts were quickly removed. Subsequently, the striatum was isolated from both hemispheres on an ice plate, stored at −80°C, and used separately for protein and gene expression studies. The isolated colons were washed in PBS and laid flat on moist paper to measure their length. After measuring their lengths, colons were -frozen and stored at −80°C for further use. The remaining half of the animals (n=20) were anesthetized using a cocktail of ketamine and xylazine, followed by transcardial perfusion initially with ice-cold phosphate buffer saline (PBS), then by 4% paraformaldehyde (PFA) in 0.05 M PBS (pH 7.4). Colons were isolated from each animal (n=5 animals per group) and fixed immediately in 4% PFA overnight. After fixation, medial colonic tissue was embedded in paraffin. The paraffinized tissue blocks were then stored at room temperature. 6μm sections were cut using the microtome and further used for histology and immunofluorescence staining. Subsequently, the brains were extracted and fixed in 4% PFA overnight., The next day, brains were then cryoprotected by immersing sequentially in 15% and 30% sucrose solution (in 0.1M PB). Furthermore, for immunohistochemical processing, coronal sections (40μm of thickness) were cut on a cryostat (Thermo Fisher Scientific). For each mouse, three coronal sections were obtained according to the following stereotaxic coordinates: A) DMV: −7.76 to −7.48mm B) LC: −5.68 to −5.34mm C) SNpc: −2.80 to −3.64mm D) Striatum: 1.34 to 1.1mm, relative to bregma, according to the mouse brain atlas^29^.

### 2.8. Hematoxylin & eosin and immunohistochemistry on colonic tissues

For Hematoxylin & eosin (H&E) staining, 6μm transverse sections were sliced from the paraffin block using a microtome (Histocore Multicut, Leica). Histologic assessment and scoring of colon tissues were carried out in a blinded fashion based on previously defined parameters. Briefly, the assessment included reporting edema, injury extent, and crypt abscesses. In this grading system, inflammation severity was scored using a scale of 0–3 (0, no inflammation; 1, slight inflammation; 2, moderate inflammation; and 3, severe inflammation), as the extent of injury (0, no injury; 1, mucosal injury; 2, mucosal and submucosal injury; and 3, transmural injury). Crypt damage was scored using a scale of 0–4 (0, no damage; 1, a basal third was damaged; 2, basal two-thirds was damaged; 3, only the surface epithelium was intact; and 4, loss of the entire crypt and epithelium). The total histopathological score was determined from the scores for each parameter to reflect the overall degree of inflammation within each specimen^30^.

For immunofluorescence staining, 6μm paraffinized colonic sections were used according to the previous protocol with little modification^31^. In brief, the slides were deparaffinized with xylene and processed by a series of alcohol dilutions, and samples were rehydrated with water for 1 minute. Antigen retrieval was performed for 10 minutes using Pepsin reagent (R2283, Sigma). Samples were washed twice with PBS containing 0.025% triton-x for 5 minutes and then blocked with protein block (ab64226, Abcam) for 20 minutes. Primary antibodies were diluted in Antibody diluent (ab6421, Abcam) and incubated with the sections overnight at 4°C. The sections were then rinsed three times in PBS containing 0.025% triton-x before being incubated for one hour at room temperature with secondary antibodies. The primary antibodies used are α-syn, Glial acidic fibrillary protein (GFAP), Zona Occludens (ZO-1), occludin, and claudin-1. These were coupled with secondary antibodies: secondary antibodies against the applicable species. Immunofluorescence sections were counterstained using DAPI (1:1000) solution for 5 minutes. Images were acquired using a confocal scanning laser microscope at a magnification of 10x, 40x (Leica TCS SP8 Microsystem). All antibodies and dilutions are listed in Tables 2 and 3.

**Table 2:** Primary antibodies used in this study.

| Antibody | Application | Dilution | Source |
| --- | --- | --- | --- |
| Claudin-1 | IF | 1:500 | Invitrogen, 71-7800 |
| Occludin | IF | 1:500 | Invitrogen, 33-1500 |
| ZO-1 | IF | 1:500 | Invitrogen, 61-7300 |
| ChAT | IF | 1:500 | Abcam, ab131097 |
| p-α-Syn | WB | 1:1000 | Abcam, ab51253 |
| GFAP | WB, IHC | 1:1000 | Abcam, ab580 |
| GAPDH | WB | 1:10000 | Abcam, ab8245 |
| TH | IHC, WB | 1:1000 | Merck, ab152 |
| α-Syn | IF | 1:1000 | Abcam, ab1903 |
| α-Syn | WB | 1:1000 | Abcam, ab212184 |

WB, Western blotting; IF, Immunofluorescence; IHC, Immunohistochemistry

**Table 3:** Secondary antibodies used in this study.

| Antibody | Application | Dilution | Source |
| --- | --- | --- | --- |
| HRP-Mouse | WB | 1:10000 | Abcam, Ab6789 |
| HRP-Rabbit | WB | 1:10000 | Abcam, Ab6721 |
| Alexa-Fluor 488<br>Goat anti-mouse IgG | IF | 1:1000 | Abcam, ab150113 |
| Alexa-Fluor 555<br>Donkey anti-goat IgG | IF | 1:1000 | Abcam, ab150134 |
| Alexa-Fluor 647<br>Goat anti-rabbit IgG | IF | 1:1000 | Abcam, ab150079 |
| Biotinylated Rabbit | IHC | 1:50 | Vector Laboratories |

### 2.9. Immunohistochemistry on brain tissues

DAB staining was performed for TH in SNpc and striatum as described previously with minor modification^32^. For immunofluorescence staining, 40 μm free-floating coronal sections of each region (DMV, LC, SNpc, and Striatum) were processed. The sections were rinsed with 0.1 M PB three times before being blocked for 20 minutes with protein block (ab64226). Primary antibodies were diluted in Antibody diluent (ab64211) and incubated with the sections overnight at 4°C. The sections were then rinsed three times in PB containing 0.025% triton-x before incubating for one hour at room temperature with secondary antibodies. The primary antibodies used were α-syn, GFAP, tyrosine hydroxylase (TH), and Choline acetyltransferase (ChAT). These antibodies were coupled with secondary antibodies against the applicable species. Sections were counterstained with DAPI after another rinse with PB mixed with triton-x. Images were acquired using a confocal scanning laser microscope at a magnification of 10x, 40x (Leica TCS SP8 Microsystem). All antibodies and dilutions are listed in Tables 2 and 3.

### 2.10. Western blotting

Frozen striatum and colon samples were thawed, homogenized in RIPA (Radioimmunoprecipitation) buffer (25mM Tris, pH 7.8, 150mM NaCl, 0.1% SDS, 0.5% Sodium deoxycholate, 1% triton-x-100, PMSF, and protease inhibitor cocktail), and centrifuged at 12,000g and 4°C for 7 min. Further, the supernatants were collected, and the protein estimation was performed using the bicinchoninic acid (BCA) method. Equal amounts of proteins loaded in each lane and resolved in 10% and 15% polyacrylamide gels based on the desired molecular weights of proteins. The detailed procedure of western blotting analysis was described previously^33^. Primary antibodies against α-syn, p-syn, TH, GFAP, and GAPDH were probed with their respective horseradish peroxidase (HRP) conjugated secondary antibodies. The expression level was quantified by densitometric analysis with the help of Image J software (NIH, USA). All antibodies and dilutions are listed in Tables 2 and 3.

### 2.11. Reverse transcriptase polymerase chain reaction (qRT-PCR)

Total RNA was isolated from the colon and striatum tissue using TRIZOL reagent (Sigma Aldrich, USA) according to the instructions. Pure purity were quantified using a Nanodrop 2000c spectrophotometer (Thermo Scientific). Extracted RNA was then reverse transcribed using an iscript cDNA synthesis kit (Bio-Rad Laboratories, USA). The diluted cDNA (1:10) was used as a template for real-time PCR (Bio-Rad CFX 96™ Real-Time system) with an SYBR^®^ Green Supermix mixture following the manufacturer’s protocol (Bio-Rad). The sequences of the primers for each gene are listed in Table 4. The quantification of each gene was carried out by the 2^−ΔΔCT^ method with 18s as endogenous control.

**Table 4:**
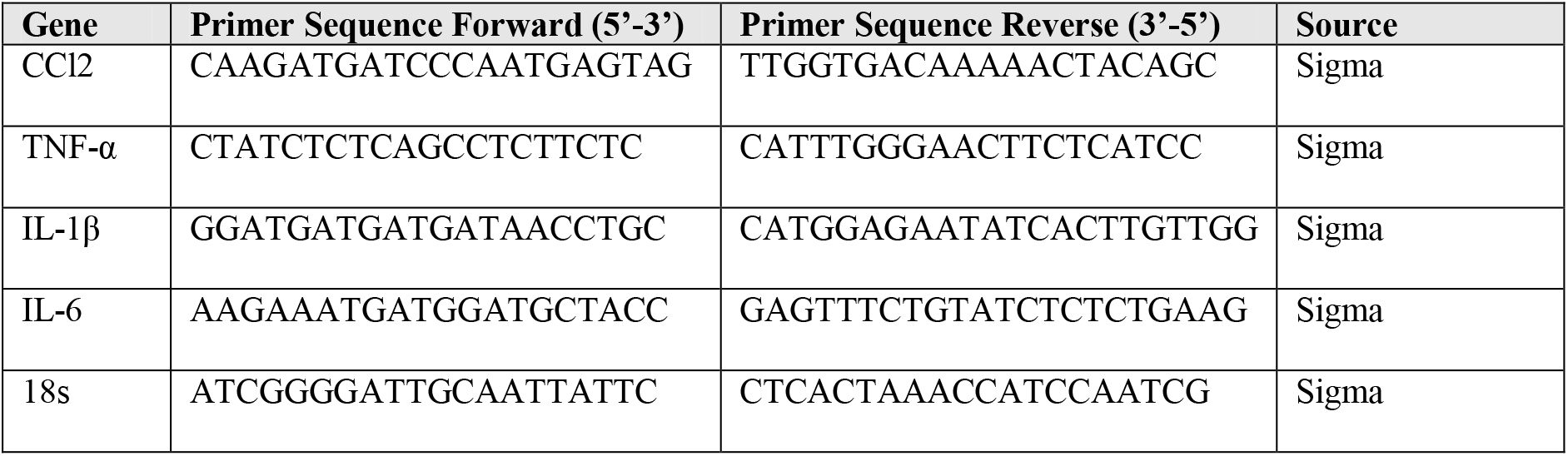
Nucleotide sequence of primers used for rt-PCR experiments

### 2.12. Fecal DNA extraction and 16S rRNA sequencing

Mice were randomly chosen from each group at the end of the 17^th^ week for microbiota sequencing. After placing each mouse in a separate empty autoclaved cage, 6-8 fresh fecal pellets were collected in a sterile tube and stored at - 80 °C for further analysis. A QIAamp PowerFecal Pro DNA Kit (Cat:51804 Qiagen, Germany) was used to extract microbial genomic DNA from feces samples based on the manufacturer’s instructions. These samples were quantified using NanoDrop One and Qubit using water and standard as controls, respectively. To check the DNA quality, all samples were aliquoted and employed on a 0.8% agarose gel in 1X Tris-acetate-EDTA buffer along with a DNA ladder, stained with Syber SAFE(Cat: S33102, Thermo Scientific) and visualized in Gel-doc (Biorad) (refer supplementary section S1). DNA was then subjected to PCR to target the 16S V3-V4 region and to generate the libraries as recommended by Illumina MiSeq sequencing (Illumina, Inc., CA, USA). Experiments, including DNA extraction, quality assessment, library construction, and high-throughput sequencing, were performed by miBiome Therapeutics (Mumbai, India).

### 2.13. 16S rRNA gene sequencing statistical analysis

Differences in alpha diversity indices of bacterial composition (Number of ASVs and Shannon index) among study groups (Control, DSS, Rot, DSS+Rot) were statistically evaluated using Kruskal-Wallis ANOVA. Beta diversity-sequencing data of bacterial abundance were treated as compositional and transformed using centered log-ratio (CLR) transformation with zerosreplaced with the constant 0.65 before all statistical analyses. Only taxa occur at least three times in relative abundance (RA) >0.1% were used for univariate and multivariate analyses. Only level of genus and level of phyla were used. Data were represented as median(min-max)

Overall gut microbiome taxonomic composition was displayed using principal component analysis (PCA), heatmaps (Euclidean distance, Ward clustering), and box plots. Univariate testing for differential abundances of each taxonomic unit among study groups was tested using Kruskal-Wallis tests corrected for multiple testing using the Benjamini-Hochberg procedure (resulting in q values). Results were considered as statistically significant for adjusted (q) value < 0.1 or possible trend for unadjusted (p) value < 0.05. All statistical analyses were performed in R, version 4.0.5^34^, using additional R packages gene filter, ver. 1.72.1 (taxa filtering)^35^, compositions, ver. 2.0.4 (CLR transformation)^36^, ggplot2, ver. 3.3.6^37^, ggpubr, ver. 0.4.0^38^, FDR estimation ver. 1.0.1 (BH procedure)^37,39,^ complexHeatmap, ver. 2.13.1 (heatmaps)^40^.

### 2.14. Statistical Analysis of protein and gene expression data

Data were represented as mean ± standard error of the mean (SEM). Two-way determined differences between groups and One-way ANOVA (analysis of variance), followed by Tukey’s multiple comparison test. Differences were considered statistically significant if p<0.05. Statistical software used for analyzing the data was GraphPad Prism, version 8.0, GraphPad Software, Inc.

#### 2.14.1. Cell counts

Images were observed, and photos were taken under a confocal microscope (Leica TCS SP8 Microsystem) as described by Ip *et al*., ^41,42.^ Six randomly selected sections over the whole anterior-posterior extent of SNpc were counted, separated by 240 μm (1/6 series). TH-immunoreactive DAergic neuronal perikarya were identified by their rounded or ovoid shape. Parameters used for TH stereological counting were as follows; counting frame size (50 μm × 50 μm) and sampling grid size (130 μm × 130 μm). The analysis was done by the authors blinded to the experimental groups. The number of ChAT^+^ neurons in the DMV was counted manually under a microscope at 40× magnification^43^.

#### 2.14.2. Optical density measurement

The extent of immunostaining of TH^+^ fibers in the striatum was presented as a percentage of optical density (OD) in control mice. Image J was first calibrated using the Rodbard function within the software to normalize the gray-scale range (0–255) into OD values. Each image was transformed into an 8-bit (grayscale). The OD values were then normalized by eliminating the OD values of the background. This was done for all three levels of the striatum and then calculated an average of these levels. The average values of rotenone mice were normalized with respect to control values^44^

The intensity of GFAP^+^ enteric glial cells (EGC) expression in the lamina propria and myenteric plexuses and α-syn expression in the myenteric plexuses of colon samples was measured using Image J software. Fluorescence images were taken at increased magnification using a confocal microscope. Images were prepared for use with Image J software, and regions of interest were chosen to quantify intensity. For each region, background measurements were obtained and removed from the mean fluorescence levels, with the corrected values shown as a percentage of control. Data were collected in the form of mean intensity measures ^31,45.^

#### 2.14.3. Integrity score measurements for tight junction proteins

A tight junction (TJ) barrier integrity score in the colon was performed as previously described ^45,46.^ Using a scale (0-3) for TJ proteins immunofluorescence sections (0 = no immunofluorescence, 1 = very light and discontinuous immunofluorescence, 2 = intense and discontinuous immunofluorescence, 3 = smooth continuous and well-organized immunofluorescence). Imaging analyses were conducted in two sections. The slides were randomized and coded before collecting fluorescence photos to avoid bias. A minimum of twenty crypts were examined, and the average data were recorded for analysis.

## 3. Results

### 3.1. Intragastric rotenone did not alter the clinical and histological aspects of DSS-induced colitis in mice but aggravated inflammatory cytokines expression

Based on the pilot study results (refer to supplementary section S2), we have selected the dose of rotenone 5mg/kg for chronic intragastric administration as this dose was below the detection threshold in both the blood and the brain and reflected environmental toxin exposure under real-world conditions. We evaluated the effect of rotenone administration on post-colitis severity. The effect of rotenone administration on DSS-induced colitis severity was evaluated by comparing the % body weight, DAI score, colon length, and histopathology among all four groups. During the development of colitis, mice began to displayed clinical signs of disease after every cycle of 1% DSS administration (weeks 1, 4, and 7), as evidenced by a significant loss in % body weight and an increase in DAI score compared to control mice (Figure 1B, 1C). The most evident clinical signs were recorded between the 6^th^ and 7^th^ weeks, with a significant weight loss that peaked with maximum DAI score, without any mouse mortality.

Compared to control and Rot alone treated mice, administration of rotenone from the 9^th^ week elicited sustained significant changes in % body weight and DAI scores in the DSS+Rot group mice. Interestingly, animals in the DSS+Rot and DSS alone groups did not restore their normal weights and had shorter colon lengths after colitis was induced (Figure 1B, 1D, 1F). Morphologically, DSS+Rot and DSS alone treated animals showed distorted crypt architecture with massive infiltration of inflammatory cells and more crypt damage than non-colitic mice (control and Rot alone group) which was then confirmed by histopathological scores(Figure 1E, 1G). Overall, these results show that, in our experimental setting, low dose of intragastric rotenone did not aggravate the histological and clinical features of colitis.

To further explore the potential contribution of rotenone administration in an IBD setting using the DSS-induced colitis mouse model. We investigated the mRNA level of inflammatory cytokines like TNF-α, IL-6, IL-1β, and CCL2 in colons samples of all four groups. Administration of 1% DSS resulted in lasting pro-inflammatory cytokine mRNA alteration in the colon. Notably, we found that there was a significantly higher level of TNF-α (*p*<0.01), IL-1β (*p*<0.05), and CCL2 (*p*<0.01) in the colon of the DSS alone group. In contrast, there were non-significant trends toward the elevation of IL-6 levels in the DSS alone group compared to the control group. Moreover, compared with rotenone alone treatment, rotenone exposure post-colitis significantly upregulated levels of inflammatory cytokines, i.e., TNF-α (*p*<0.001), IL-6 (*p*<0.01), IL-1β (*p*<0.001) and CCL2 (*p*<0.001) in colon respectively (Figure 1H). Interestingly, we also found the elevated expression of TNF-α and CCL2 in the colon of Rot alone group mice compared to the control.

### 3.2. Intragastric rotenone post-colitis aggravates gastrointestinal dysfunction and accelerates the onset of motor impairment without eliciting extraintestinal inflammation in colitis mice

To check whether alterations at the intestinal level were linked with the functional outcome of PD, we evaluated fecal pellet output and fecal water content at 0, 9^th^, 13^th^, and 17^th^ weeks of our study. With respect to normal initial status (0, baseline), there was no noticeable difference in water reabsorption in the colon between the groups, as assessed by fecal water content. Furthermore, from the 13^th^ to the 17^th^ week, DSS+Rot group mice had significantly lower fecal pellet counts (*p*<0.01) and percent water content (*p*<0.05) than control mice, which is to be expected given that constipation is a common early symptom of PD (Figure 2A, 2B).

**Figure 2:**
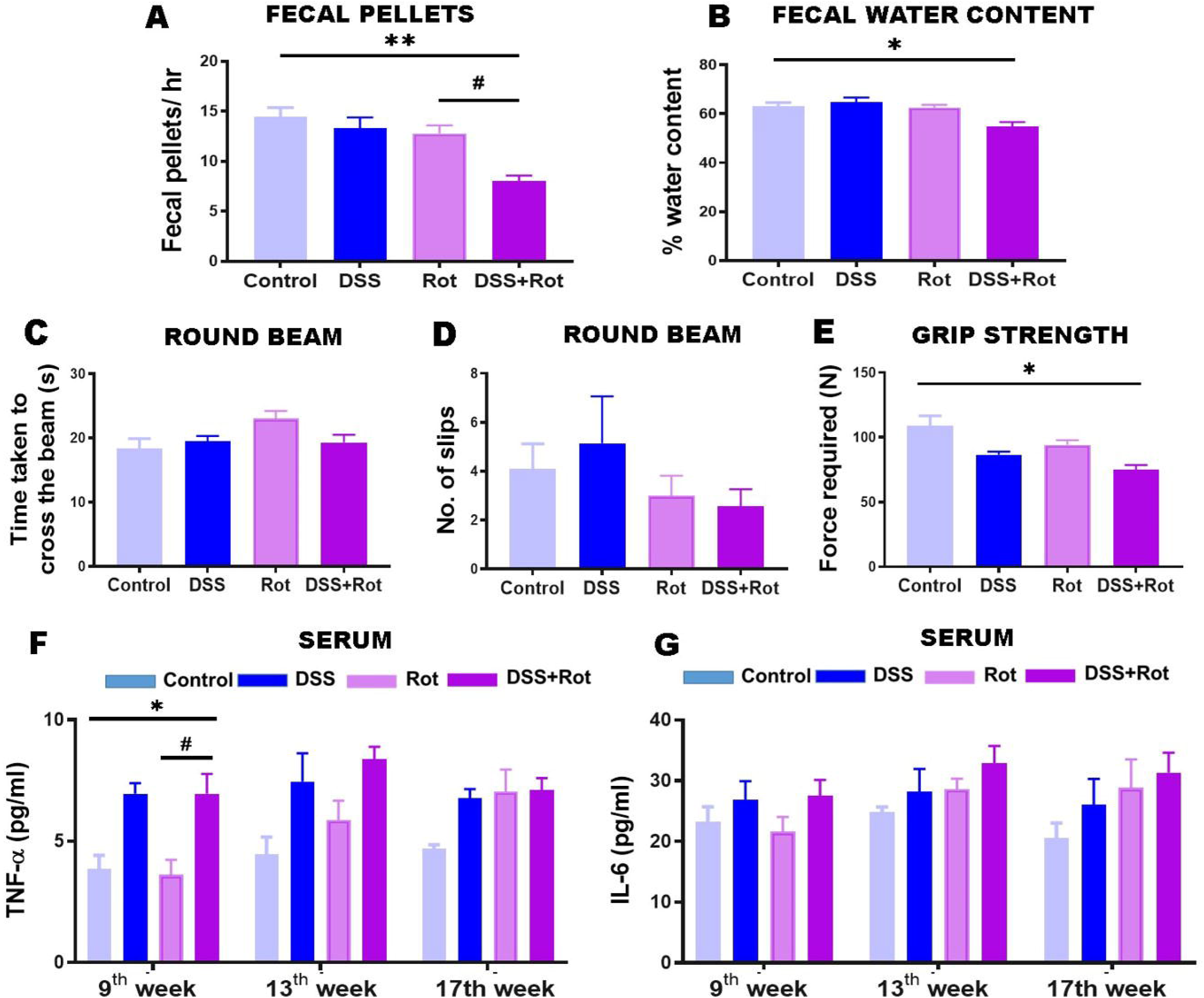
A) The numbers of total fecal pellets during 1 hr. fecal weight (B) and water content gradually decrease with the extension of the rotenone treatment time. Effect of a low dose of rotenone on round beam walk. C) Bar plot represent the time taken to cross the beam D) Bar plot represents the number of slips. E) Bar plot represent force required on grip strength is significantly lower in the DSS+Rot group compared to the control group(n=8-12). Effect of a low dose of rotenone on Serum inflammatory cytokines G) TNF-α, and H) IL-6 in Serum was measured by ELISA at 9^th^, 13^th^, and 17^th^ weeks (n=4-5). Data were expressed as Mean ± SEM. *p< 0.05 **p< 0.01 vs. control. ^#^p< 0.05 vs. rotenone. Statistical analysis was performed using one and two-way analysis of variance (ANOVA) followed by Tukey’s multiple comparison test.

A general evaluation of the motor activity of mice chronically exposed to rotenone post-colitis was performed using the RBW and grip strength test. Any changes in locomotor characteristics measured by the RBW and the grip test were detected at 0, 9^th^, 13^th^, and 17^th^ weeks during rotenone administration on DSS-challenged mice. During the establishment of the PD model post-colitis, we observed that chronic rotenone administration exhibited no significant changes in the time taken to cross the beam and no slips in DSS+Rot animal group compared to control group at any time point. Whereas, at the 17^th^ week, when the nigrostriatal degeneration was maximal, a strong trend was observed in the time taken to cross the beam on the DSS+Rot group relative to control mice (Figure 2C, 2D). Moreover, we also found a significant reduction in force required to hold the grid in DSS+Rot mice when compared to the control mice (*p*<0.05) in the 17^th^ week of our investigation (Figure 2E). As anticipated, at any time point, there were no differences between the Rot alone group and the control group in terms of the time required to cross the beam, the number of slips, and the performance of the limb muscular force in the grip strength test (refer supplementary section S3 for more details). These findings revealed that rotenone administration post-colitis induces GI dysfunction before significant motor abnormalities.

To evaluate whether rotenone administration post-colitis led to extraintestinal inflammation, we measured the inflammatory cytokines (TNF-α, IL-6) in Serum at different time points. DSS treatment significantly elevated the contents of pro-inflammatory TNF-α *(p*<0.05) level before the administration of rotenone, which was further normalized in a subsequent week (Figure 2F). Moreover, we could not find any significant changes in serum IL-6 levels at any time point (Figure 2G). These results revealed that chronic colitis induced a significant elevation in the gene expression of the colon’s pro-inflammatory marker, resulting in the initial detection of TNF-α in the blood (Figure 2F).

### 3.3. Chronic exposure to intragastric rotenone post-colitis promoted intestinal tight junction disruption, α-syn accumulation, and GFAP levels in colonic myenteric Plexus in Mice

Compromised intestinal barrier function is hypothesized to promote intestinal inflammation and systemic peripheral inflammation, a phenotype in which PD and IBD share similarities. Herein, to evaluate intragastric rotenone’s effect on the intestinal barrier function of colitis mice, we checked the expression level of three key junction proteins, i.e., ZO-1, occludin, and claudin-1 by immunofluorescence. Immunofluorescence staining showed robust expression of ZO-1 at the colonic epithelial lining of control and Rot alone-treated mice. However, administration of intragastric rotenone post-colitis disrupted and decreased expression of ZO-1 at the epithelial lining of colitis mice, irrespective of the inflamed mucosa. Semi-quantitative analysis of staining using an integrity score scale (0 = no staining – 3 = continuous normal expression) showed a significant difference due to rotenone exposure in DSS+Rot mice *(p*<0.01), but a non-significant decrease was observed in DSS alone treated mice (Figure 3A, 3B). In the case of occludin and claudin-1, microscopic visualization of immunofluorescence labeled junction protein demonstrated robust expression of both proteins at the colonic epithelial lining of the control and Rot group (Figure 3A, 3C, 3D). On the contrary, semi-quantitative analysis of staining using integrity scale score revealed a significant decrease in expression of these junction proteins in DSS alone treated mice, and rotenone exposure further reduced the level of proteins *(p*<0.01, *p*<0.001 respectively). These results indicate that expression and localization of TJ proteins were inhibited by long-term rotenone exposure after colitis.

**Figure 3:**
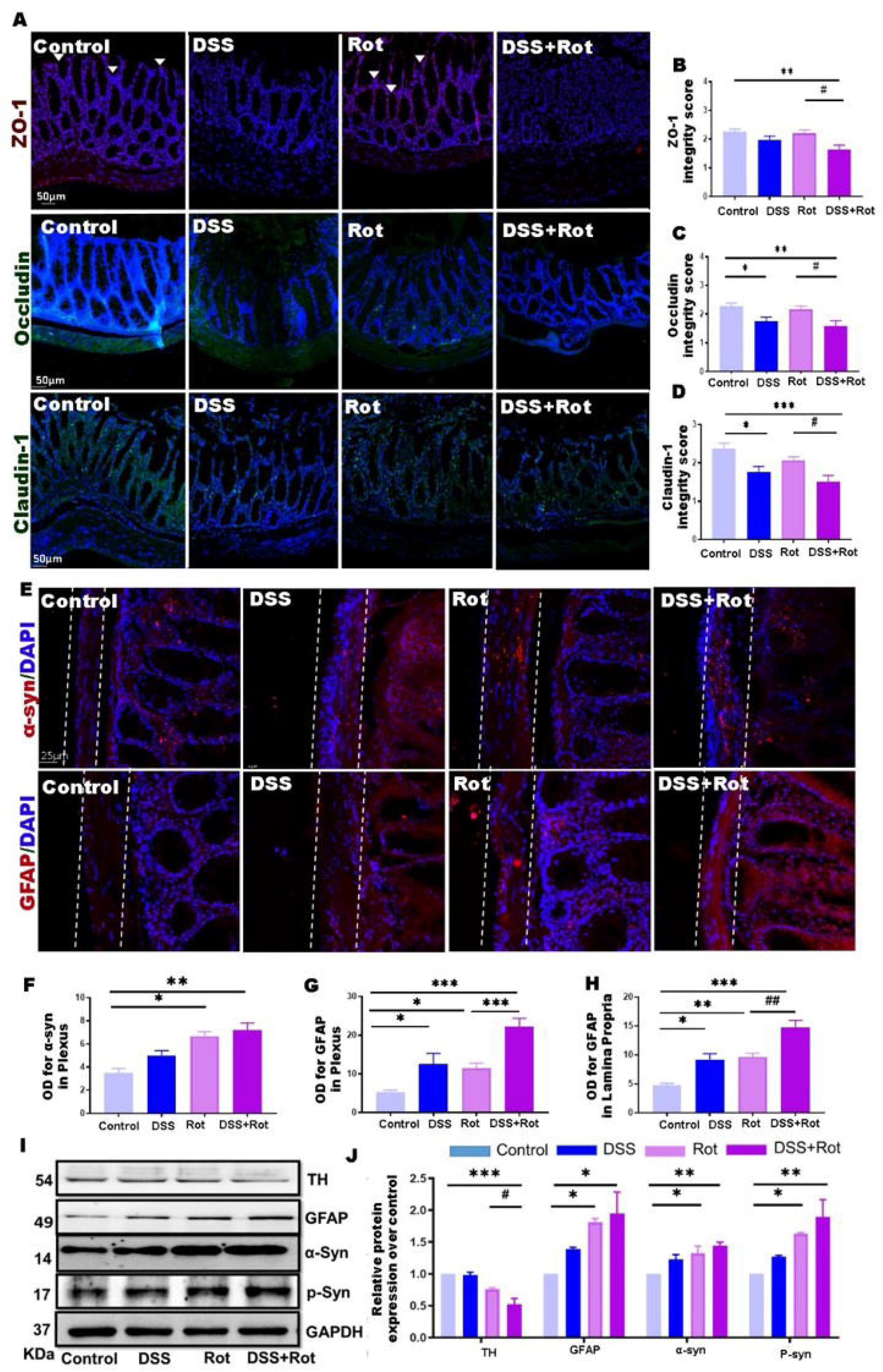
A) Representative immunofluorescence images for the tight junction proteins ZO-1, occludin, and claudin-1 in the mouse colonic epithelial cells. Arrows in Panel ZO-1 show the red staining of ZO-1 protein on the top of the intestinal crypts. An arbitrary scale of 0–3 scores (0 = no expression - 3 = continuous normal expression of the barrier) was used to score barrier integrity. B) Integrity scoring box plot for ZO-1, (C) integrity scoring box plot for occludin, and (D) Integrity scoring box plot for claudin-1. E) Representative images of the expression of α-syn and enteric glial cells (GFAP+) in the myenteric plexuses. (F) Bar plot represents optical density analysis for α-syn expression in the myenteric plexuses (G, H) show OD analysis of GFAP expression in the myenteric plexuses and lamina propria I, J) Representative western blot image and quantification bar plot showing the effect of low dose administration of rotenone on protein expression of, TH, GFAP, α-syn, p-syn in the colon of chronic colitis mouse model (n=4-5). Data were expressed as Mean ± SEM. *p< 0.05 **p< 0.01 vs. control. ^#^p< 0.05, ^##^ p< 0.01 vs. rotenone. Statistical analysis was performed using one-way analysis of variance (ANOVA) followed by Tukey’s multiple comparison test. The scale bar in A and E represent 50 and 25 μm, respectively. OD = optical density, ZO-1= Zona occludens-1, GFAP = Glial fibrillary acidic protein, TH= Tyrosine hydroxylase, α-syn = alpha-synuclein.

Next, we assessed intragastric rotenone’s effect onPD-like pathology (specifically α-syn structures) in the myenteric plexus of an inflamed colon. Optical density analysis in myenteric plexus revealed that intragastric rotenone for 8 weekss significantly induced colonic expression of α-syn structures (p<0.05) in Rot alone mice compared to control mice. The presence of preexisting colonic inflammation further aggravated the α-syn expression in the colonic myenteric plexus of DSS+Rot (p<0.01) compared to the control group. We also observed the trend toward increased α-syn expression in the myenteric plexus of the DSS alone group compared with the control group (Figure 3E, 3F). Immunofluorescence data were further confirmed by western blot analysis of colonic tissue, showing a significantly higher expression of α-syn in the colonic tissue of Rot alone groups. However, combined treatment of rotenone post-colitis magnifies α-syn overexpression *(p*<0.01) in the colon. We also observed significant hyperphosphorylation of α-syn (p-syn) in colonic tissue of DSS+Rot as well as the Rot alone group *(p*<0.01, *p*<0.05), but no effect was observed on the p-syn expression of DSS alone mice (Figure 3I, 3J). These results suggest that rotenone exposure, irrespective of colitis, significantly exacerbated the expression of α-syn expression and its phosphorylation in the colon.

Our next step was to determine if immune marker changes accompany these changes. Thus, we examined the GFAP as a marker of enteric glial reactivity in lamina propria and myenteric plexus of colon tissue. While Rot and DSS alone showed an increased expression/number of GFAP^+^ enteric glial in myenteric plexuses and lamina propria compared with the control group. In contrast, to control mice, rotenone administration after colitis increased the expression of GFAP^+^ enteric glial cells in myenteric plexuses and the lamina propria. Post hoc analysis showed that GFAP intensity in the myenteric plexus of DSS+Rot compared to controls *(P*<0.001), Rot compared to controls (*P*<0.05), and DSS+Rot compared to Rot alone *(P*<0.001). Western blot analysis confirmed these findings regarding the relatively higher expression of GFAP+ reactivity. Our results show a significantly higher protein expression of GFAP in DSS+Rot and Rot alone mice than in control. Whereas a substantial trend was noticed in DSS alone treated animals.

### 3.4. Intragastric rotenone in combination with colitis-induced cholinergic degeneration, Astrocyte activation, and α-syn accumulation in the DMV

It has been reported that cholinergic neurons tend to be lost in DMV’s of PD patients in the early stages^47^. Thus, we sought to examine if the local effect of rotenone on the ENS of an inflamed colon aided in the acceleration and development of α-syn pathology *via* connecting structures like the DMV. Treatment with a low dose intragastric rotenone reduced the numbers of ChAT^+^ signals in the DMV, and the presence of pre-existed colonic inflammation further reduced the number of ChAT^+^ signals (*p*<0.001) when compared to control (Figure 4A,4B). However, there was no difference in the number of cholinergic neurons per section between the control and DSS alone group. The ChAT^+^ signals in the DMV of DSS alone animals were nearly identical to those in the control group. We also found that after 8 weeks of rotenone treatment, regardless of the existence of DSS-induced inflammation, there was a significant increase in intracellular α-syn accumulation inside ChAT^+^ cells (Rot, *p*<0.05; DSS+Rot, *p*<0.01) compared to control group (Figure 4A,4C). To check if comparable astrocyte activation occurred in DMV due to intracellular α-syn accumulation, we carried out immunochemical staining of DMV samples for GFAP. Quantitative analysis showed a significant increase in astrocyte GFAP expression in DMV of Rot and rotenone-treated post-colitis (Rot, *p*<0.01; DSS+Rot, *p*<0.01) compared to control, suggesting that chronic rotenone administration itself is sufficient to induce α-syn accumulation in DMV (Figure 4D,4E).

**Figure 4:**
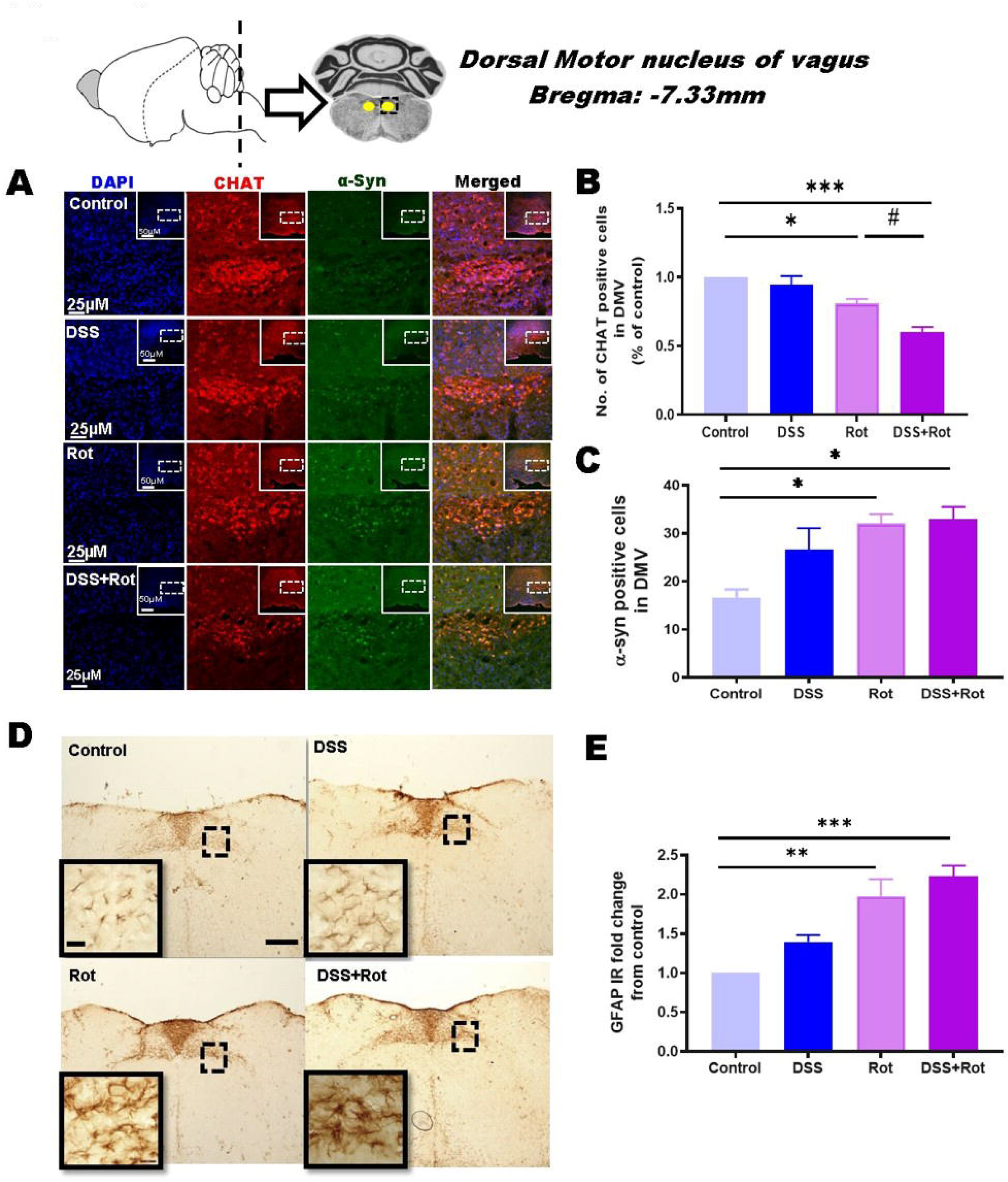
Chronic exposure to low-dose rotenone after colitis-induced cholinergic neurodegeneration and α-syn accumulation in the dorsal motor nucleus of the vagus (DMV) of mice A) Representative immunofluorescence images of DMV sections stained with rabbit anti-alpha-synuclein (green) and goat anti-ChAT antibodies (red) and with DAPI (blue) obtained all four groups (n=4-5). Scale bar in A represents 50 and 25 μm B) Bar plot represents changes in the number of ChAT+ neurons inside the DMV C) Bar plot represents the number of α-Syn^+^ cells in DMV in all four groups. D) Representative DAB images of GFAP expression in DMV of all four groups. Magnification: 10X and 40X; Scale bar: 500 and 25μm E) Bar plot represents the mean fluorescence intensity of GFAP in all four groups. Data were expressed as Mean ± SEM. (n=4-5) *p< 0.05 **p< 0.01, ***p< 0.001 vs. control. ^#^p< 0.05, ^##^p< 0.01 vs. rotenone. Statistical analysis was performed using one-way analysis of variance (ANOVA) followed by Tukey’s multiple comparison test. The scale bar in A represents 50 and 25 μm.

### 3.5. Intragastric rotenone in combination with colitis showed loss of tyrosine hydroxylase neurons with subsequent α-syn accumulation and astrogliosis in the locus coeruleus

As discussed above, α-syn pathology does not develop simultaneously in all vulnerable brain regions but in a sequential way. Both clinical and imaging studies, as well as neuropathological evidence, show that the region of the noradrenergic LC system is involved early in the topographical sequence of pathological changes in rapid eye movement sleep behavior disorder (RBD), a specific prodromal stage of PD, years before the DAergic SNpc is affected and motor symptoms become apparent in PD^48,49.^ To determine whether TH expression in LC is affected by the rotenone treatment in conjunction with chronic colitis, we assessed TH-IR in a section from all four groups. After 8 weeks of intragastric rotenone, Rot alone treated animals showed a non-significant increase in TH-IR compared to the control. On the contrary, DSS alone mice showed a strong trend of decreased TH expression compared to control mice. Furthermore, a combination of rotenone after induction of colitis significantly aggravated TH-IR downregulation (*p*<0.05) as compared to control mice (Figure 5A, 5B).

**Figure 5:**
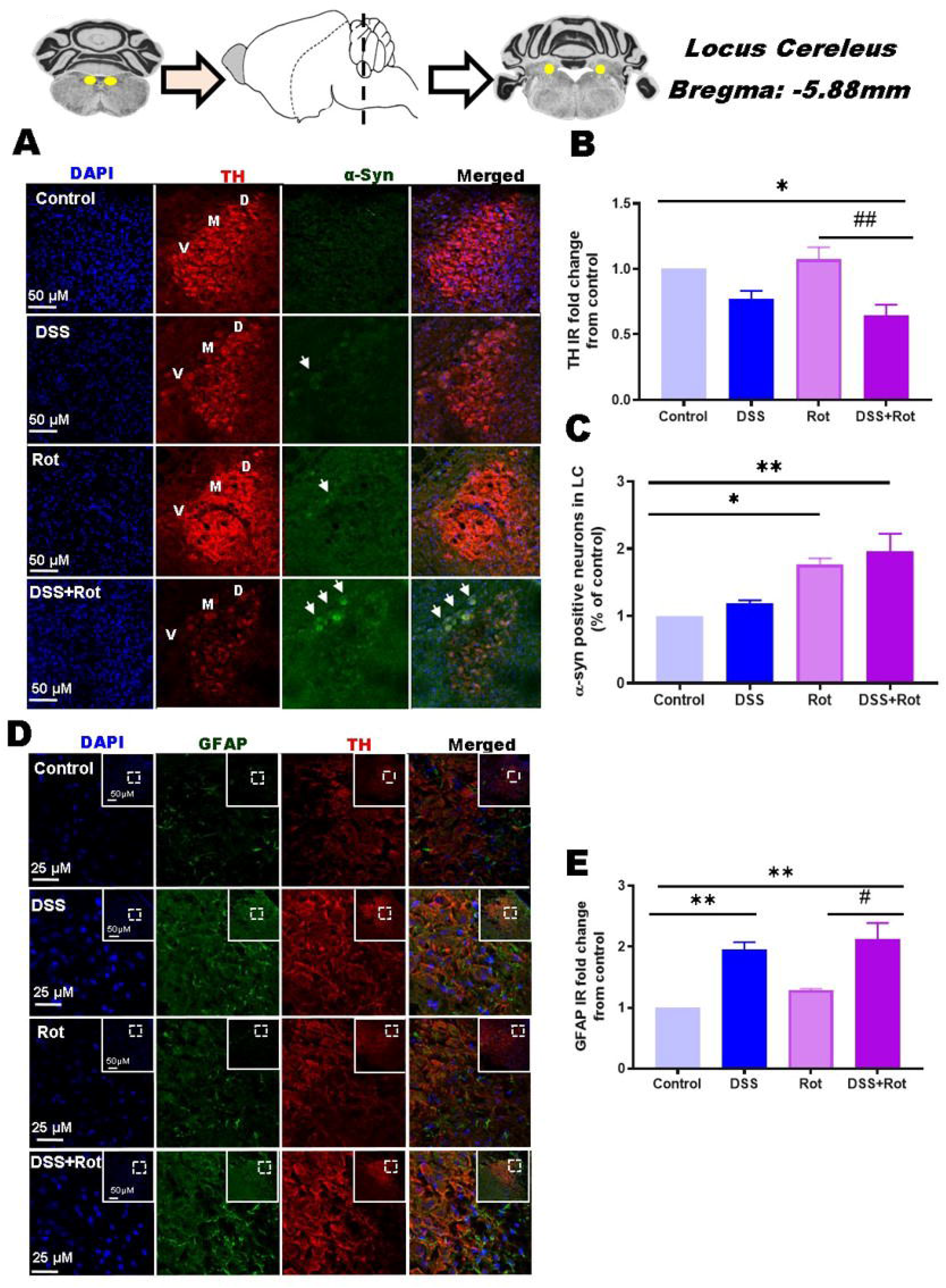
A) Representative immunofluorescence images of α-syn and TH expression in LC of mice from different groups. B) Bar plot represents the TH^+^ intensity in LC (percentage of control). C) Bar plot represents the number of α-syn^+^ neurons (percentage of control) in the LC compared between different groups. D) Representative images obtained after immunofluorescence staining for GFAP and TH expression in LC of mice from different groups. E) Bar plot represents GFAP immunoreactivity fold change compared between different groups. The results were expressed as Mean ± SEM (n = 4-5). * p < 0.05, ** p < 0.01, vs. control group, ^#^p< 0.05, ^##^p< 0.01 vs. rotenone. Statistical analysis was performed using one-way analysis of variance (ANOVA) followed by Tukey’s test.

Next, we analyzed α-syn protein in the LC by immunofluorescence. Using an antibody specific for α-syn, the expression of α-syn pathology was found to co-localize with TH-expressing LC neurons only in the sections from colitis mice (DSS+Rot, DSS) regardless of rotenone treatment (Figure 5A). However, there was a significant increase in α-syn positive neurons *(p*<0.01) in LC of DSS+Rot mice compared to control mice. On the other hand, no α-syn specific immunofluorescence was not detected in LC neurons of control and Rot alone mice (Figure 5A, 5C).

Many studies suggest that dysregulated DAergic neurotransmission is associated with inflammation and that deregulation could contribute to chronic neuroinflammation observed in PD^50^. To determine if comparable astrocyte activation occurred in LC, we carried out immunochemical staining of LC samples for GFAP. Quantitative analysis showed a significant increase in astrocyte GFAP expression in the dorsal part of LC of colitis mice irrespective of rotenone treatment (DSS+Rot, *p*<0.001; DSS, *p*<0.001) compared to control. GFAP^+^ staining was observed around TH^+^ stained neurons in the LC but not inside the LC (Figure 5D, 5E). These results suggest that α-syn pathology was restricted to DMV in the rotenone alone group, causing degeneration of ChAT^+^ neurons. In the DSS+Rot group, α-syn pathology progressed to LC, causing degeneration of TH^+^ cells accompanied by elevated GFAP expression.

### 3.6. Intragastric rotenone-induced a-syn accumulation, astroglia activation, and loss of DAergic neuronal death in SNpc of a colitis-challenged mouse

To explore the progression of α-syn pathology from the brain stem to SNpc, we have performed double immunostaining for α-syn and TH in the midbrain coronal section. Quantitative analysis revealed that there was a significant increase in α-syn mean fluorescence intensity in TH^+^ neurons of DSS+Rot mice (*p*<0.01) compared to the control group (Figure 6A,6B). It is well known that the accumulation of α-syn contributed to the neuropathological degeneration of TH^+^ neurons in SNpc. Next, we performed DAB staining on coronal sections from three levels of SNpc and counted the TH^+^ neurons as per the previously described method^51^. We found a significant decrease in the TH+ neurons in the DSS+Rot group (*p*<0.001) compared to control animals. No significant reduction was observed in the Rot and DSS alone group compared to the control group (Figure 6D, 6F).

**Figure 6:**
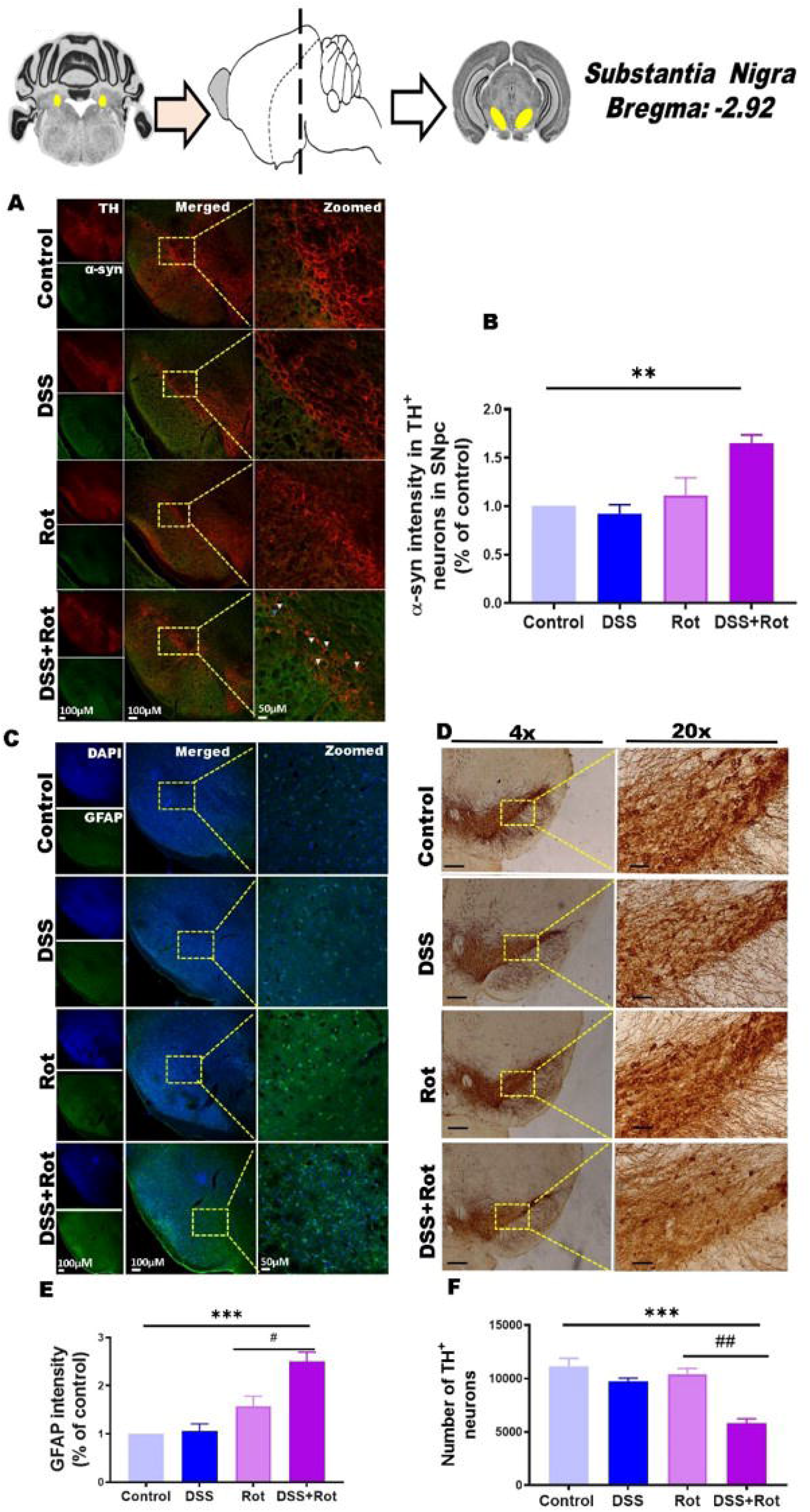
Representative immunofluorescence images of α-syn and TH expression in SNpc of mice from different groups. B) Bar plot represents α-syn intensity (percentage of control) in the SNpc compared between different groups. C, E) Representative immunofluorescence images of GFAP in SNpc of mice from different groups. The bar plot represents GFAP mean fluorescence intensity (percentage of control) in the SNpc compared between different groups. D, F) Representative immunofluorescence images of TH^+^ cells in the SNpc at 4x and 20x (scale bar: 200 μm and 50 μm, respectively). The bar plot represents the number of TH^+^ cells in SNpc. The results were expressed as Mean ± SEM (n =4-5). ** p < 0.01*** p < 0.001, vs. control group, ^#^p< 0.05, ^##^p< 0.01 vs. rotenone. Statistical analysis was performed using one-way analysis of variance (ANOVA) followed by Tukey’s test.

It has been reported that astrocyte activation may contribute to neuroinflammation and DAergic neuronal death in SNpc during PD development^52^. In our present study, we performed immunofluorescence staining of the SNpc regions to detect the activation of glial cells, using GFAP as a marker of gliosis. We found a significant elevation of GFAP^+^ cells intensity in the SNpc of DSS+Rot mice compared with control (*p*<0.001) and Rot alone (*p*<0.05) treated mice (Figure 6C,6E). While preexposure to DSS in mice showed a non-significant increase in GFAP intensity compared to the control group. These results suggest that low-dose rotenone administration after colitis caused accumulation of α-syn, astrogliosis resulting in degeneration of DAergic neurons in SNpc.

### 3.7. Intragastric rotenone-induced a-syn accumulation, astroglia activation, and loss of DAergic fibers in Striatum of colitis-challenged mouse

Once the α-syn pathology reaches SNpc, we wanted to see its progression and effect on TH^+^ fiber in the striatum. Quantitative analysis revealed that chronic treatment with rotenone after colitis induced a significant reduction in the density of striatal TH^+^ fibers (*p*<0.001) compared to the control. On the other hand, mice did not show any significant changes in the density of striatal TH^+^ fibers when exposed to rotenone or DSS alone (Figure 7A, 7C). Next, we performed a western blot to check the protein expression of p-syn and GFAP in the striatum. Here, we found a significant increase in p-syn levels (p<0.05) in the DSS+Rot group relative to control mice. As well, GFAP expression is also significantly increased in the DSS+Rot group (*p*<0.001) as compared to the control group (Figure 7E, 7F, 7G). This also indicated that rotenone alone could not change p-syn and GFAP expression in the striatum at 8 weeks of administration. However, the presence of existing inflammation aggravated the p-syn and GFAP overexpression. Meanwhile, we also examined the mRNA levels of TNF-α, IL-6, IL-1β, and CCL2 in the striatum of all four groups. Pro-inflammatory cytokine mRNA was altered in the striatum in a long-lasting way after administration of rotenone after 1% DSS. The levels of TNF-α (*p*<0.001), IL-6 *(p*<0.001), IL-1β (*p*<0.001), and CCL2 (*p*<0.01) were significantly higher in the striatum of DSS+Rot compared to control group. In addition, the levels of TNF-α, IL-6, IL-1β, and CCL2 in the striatum were not altered in DSS mice compared with control (Figure 7H, 7I, 7J, 7K).

**Figure 7:**
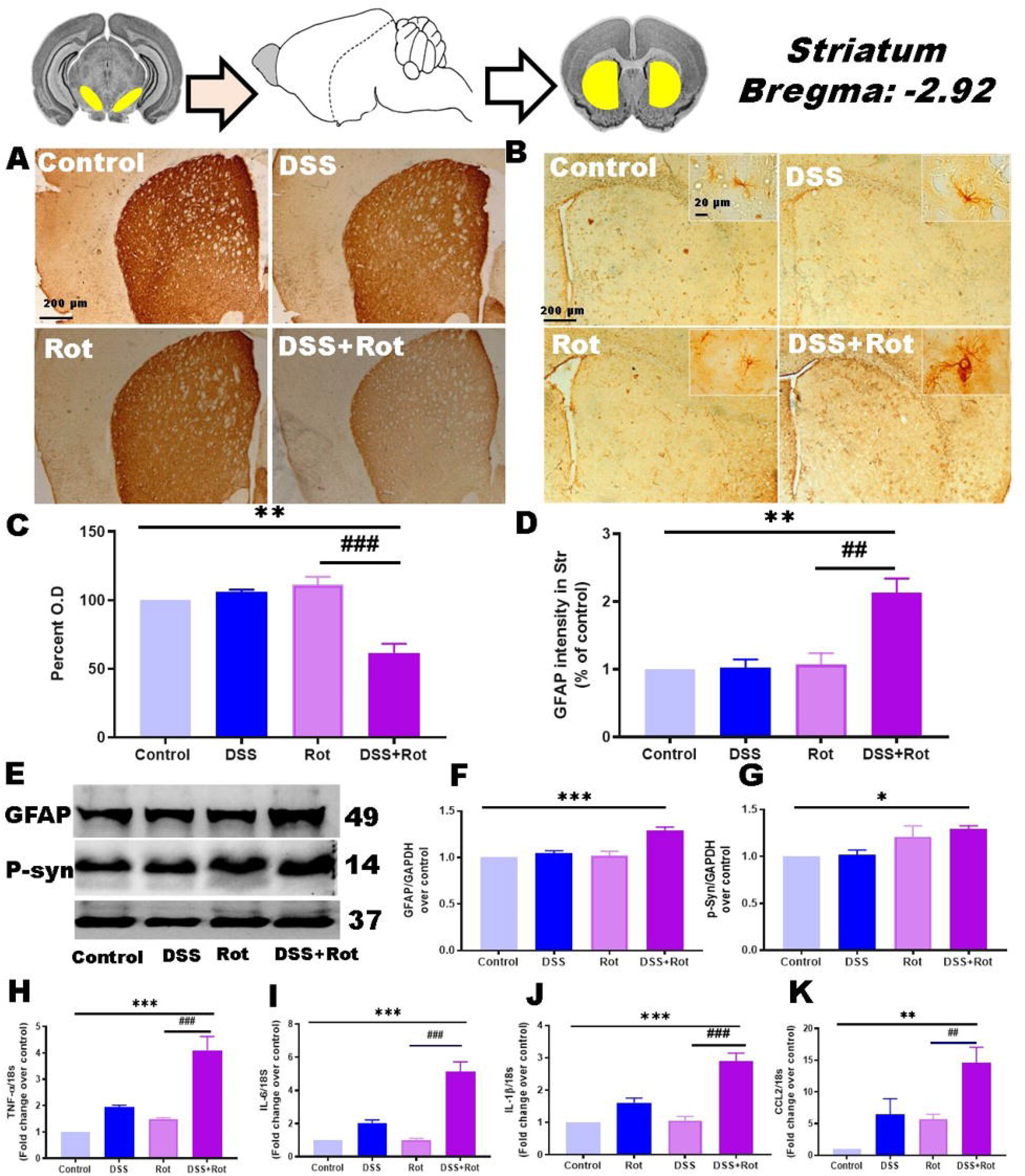
Representative immunofluorescence images of TH^+^ fibers in the striatum at 4X (scale bar: 200μm). B) Representative immunofluorescence images showing the effect of low-dose administration of rotenone on GFAP protein expression. C) Bar plot represents the mean optical density (OD) of TH^+^ fibers in the striatum. Values of OD are expressed to 100% for levels of the control group. D) Bar plot represents the mean O.D of GFAP expression in the striatum. Values of OD are expressed to 100% for levels of the control group. E) Representative western blot image and quantification bar plot showing the effect of low-dose administration of rotenone on protein expression of (F) GFAP and (G) p-Syn in the striatum of a chronic colitis mouse model. H, I, J, K) Relative gene expression levels of TNF-α, IL-6, IL-1β, and CCl2 in striatum from all four groups measured using RT-PCR, normalized to housekeeping gene 18s, and presented as fold change relative to controls. The results were expressed as Mean ± SEM (n = 5). * p < 0.05, ** p < 0.01, ***p<0.001 vs. control group, ^#^p < 0.05, ^##^p < 0.01, ^###^p<0.001, vs. rotenone group. Statistical analysis was performed using one-way analysis of variance (ANOVA) followed by Tukey’s test.

### 3.8. Disruptions in the gut microbiome due to low-dose rotenone exposure post-colitis

We investigated how low-dose rotenone affected the gut microbiota composition in DSS-induced microbial dysbiosis. We performed 16S rRNA amplicon-based sequencing to assess the composition of gut microbes. Simpson and Shannon diversity indices showed no differences between groups (data not shown), suggesting that the alpha diversity of gut microbiota was not impacted by DSS or Rot administration. Moreover, PCA results showed that intestinal microbiome clusters in all three mice groups (DSS, Rot, and DSS+Rot) dramatically differed from those of the age-matched control mice (Figure 8A). Notably, the phylum *Firmicutes/Bacteroidetes* (F/B) ratio showed a decreasing trend t (p < 0.05, q > 0.1) in DSS alone and DSS+Rot mice compared with control mice (Figure 8C). Compared to the control group, we found a lower relative abundance of *Lactobacillus* at the genus level in the DSS alone and DSS+Rot groups (Figure 8D).

**Figure 8:**
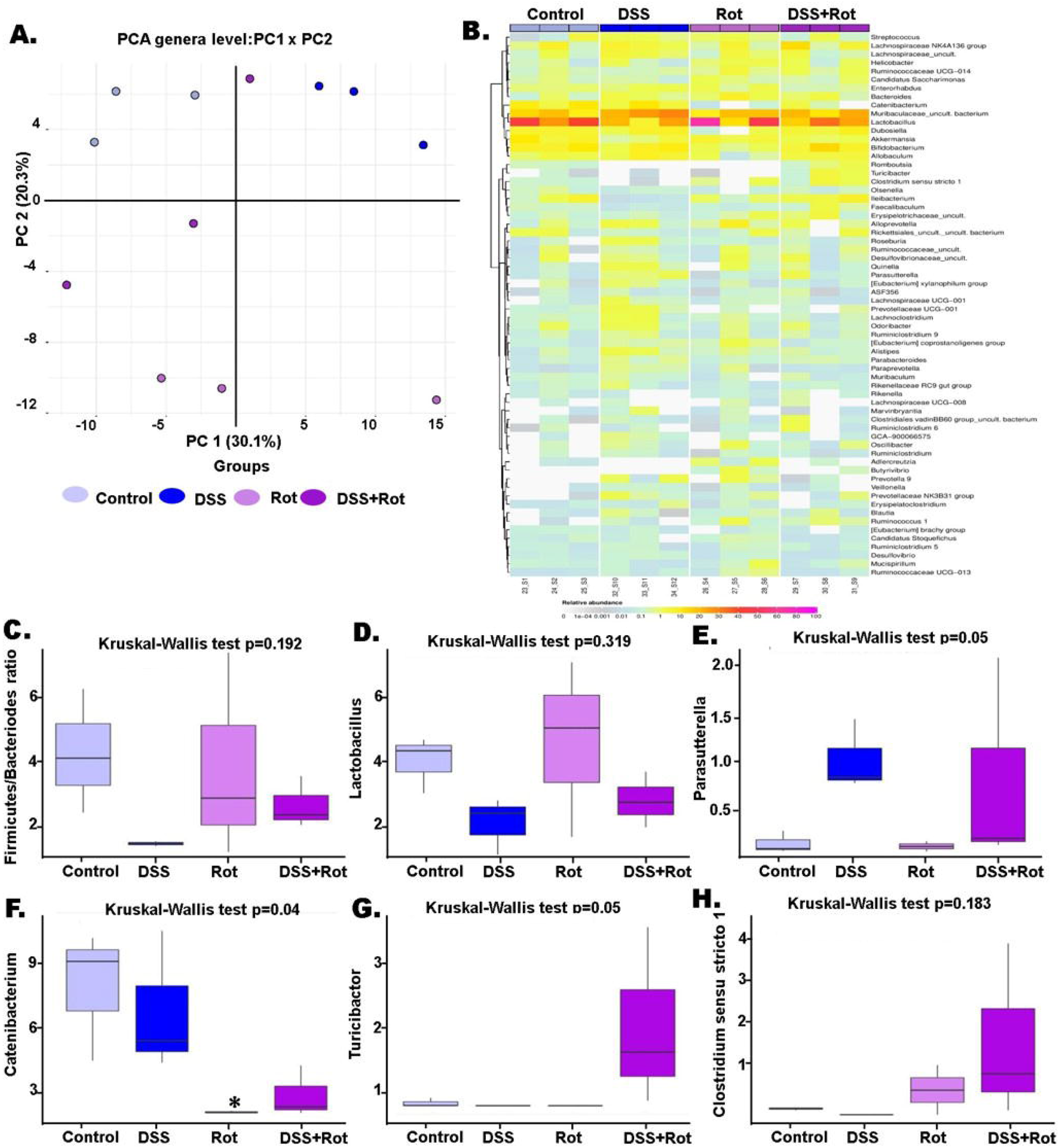
Changes in gut microbiota composition following rotenone or DSS treatment in mice (A) PCA plot for the four experimental groups of interest (control, DSS, Rot, and DSS+Rot). (B) Heatmap of microbial composition at genera level, the color intensity shows RA (C) Box plot represents the ratio of Firmicutes to Bacteroidetes (D) Box plot represents the relative abundance of Lactobacillus (E) Box plot represents the relative abundance of Parasutterella; (F) Box plot represents the relative abundance of Catenibacterium (G) Box plot represents the relative abundances of Turicibacter H) Box plot represents the relative abundance of Clostridium sensu stricto 1. (E, F, and G) show a possible trend (Kruskal-Wallis ANOVA unadjusted p < 0.05 and no statistically significant result for post hoc Dunn test).

Furthermore, at the genus level, *Parasutterella* has significantly increased in DSS alone mice, with an elevated level in the DSS+Rot group compared to the control group that can be positively correlated with chronic intestinal inflammation and the severity of PD (Figure 8E). We also found that rotenone treatment after colitis decreased the RA of the bacterial genus *Catenibacterium* (Figure 8F). There were changes in the RA of two bacteria genera after DSS and rotenone treatment, such as increased *Turicibacter* and *Clostridium sensu stricto 1*, which were not seen following DSS alone treatment when compared to the control group (Figure 8G, 8H). Interestingly, the microbial changes we observed in our DSS-alone treated mice, such as increased *Parasutterella* and decreased *Catenibacterium*, have been linked to worsening clinical symptoms in human IBD studies. There were no statistically significant results in differential abundance testing after adjustment for multiple comparisons (due to small sample size), all of the described results were significant unadjusted (p < 0.05).

## 4. Discussion

A growing body of preclinical and clinical research suggests that GI inflammation integrates α-syn pathologic alterations and progressive neurodegeneration in driving chronic PD progression. Globalization’s impact has subjectedthe world’s population to a staggering risk of IBD, as represented by around 10 million people worldwide. Importantly, recent meta-analysis reports suggest that patients with the onset of IBD over 60 have a 32% higher risk of PD than those without IBD^12,53.^ However, it is still to be determined if IBD patients are susceptible to developing PD because of a higher inflammatory milieu in the GI tract or if there are some other genetic or environmental factors along with GI inflammation that make them vulnerable to developing PD. Researchers have used transgenic (Tg) mice either overexpressing α-syn or susceptible genes that are dysregulated in PD to test the effect of DSS-induced colitis on α-syn accumulation and subsequent DAergic neurodegeneration^17,54^

Moreover, it will be interesting to study the interaction of IBD with environmental toxins, such as exposure to pesticides, as the world is facing an unprecedented crisis of environmental health risks related to the excessive and unsafe use of pesticides. Consequently, it gives rise to a higher concentration of pesticide residue in the food chain, as evidenced by overall usage, especially in the food industry and agricultural sectors. This provides an opportunity to investigate potential aetiopathogenetic factors that may result in the onset of PD pathologies, such as α-syn accumulation in the GI tract and subsequent progression to the brain via the DMV. As the incidence of IBD and long-term pesticide exposure through food rises with age, it is crucial to comprehend how early colitis among patients influences the probability of other neurodegenerative diseases such as PD. This work aimed to investigate the susceptibility of IBD patients in the presence of environmental toxins to develop PD using DSS-induced colitis and progressive intragastric rotenone mouse model. More specifically, how early GI (colonic) inflammatory milieu interacts with environmental toxins to mediate α-syn pathology and worsen the PD phenotype in a progressive intragastric rotenone mouse PD model.

To achieve this, we first optimized the concentration of DSS that is adequate to cause moderate GI inflammation with alterations in the expression of TJ proteins only in the colon but not in the brain tissue of mice (refer to supplementary section S4 for more details). Previous reports also revealed that low-dose intragastric rotenone treatment for 2 months elevated the level of α-syn expression in the ENS, along with a pro-inflammatory GI state, without demonstrating rotenone presence in blood or brain^20^. As a second step, we employed the LC-MS method to detect the rotenone level in the blood and brain of early colitis mice and found an undetected level of rotenone with 5mg/kg dose in the blood or the brain (refer supplementary section S2). These two preliminary findings led to a uniform assumption that the association of 1% DSS and intragastric rotenone dose at 5mg/kg fall below the injury threshold, leaving brain TJ’s structurally intact.

The proposed and widely accepted mechanism by which DSS causes colonic inflammation involves the alteration in TJ’s protein expressions on colonic epithelial monolayer lining and pro-inflammatory changes in the GI microbiota^55,56.^ Therefore, we assessed key TJ proteins (ZO-1, Occludin, and Claudin-1) in colonic tissue. Our intestinal permeability results observed in this investigation revealed that low-dose rotenone administration for 8 weeks post-colitis caused significant permeability changes compared to the control group. Whereas no difference was observed in the integrity score of all three TJ proteins in the Rot alone group compared to the control group. It is not surprising as lower rotenone dose does not affect TJ protein expression^45^. In contrast, studies with higher rotenone doses (30mg/kg) were reported to alter all three TJ proteins mainly by altering the gut microbes^57^. Notably, we could not observe any changes in the ZO-1 integrity score in the DSS alone group, with a subsequent significant reduction in the integrity score of occludin and claudin-1 compared to the control group. This indicated that disruption of these TJ proteins represents an adaptive response to the toxin or changes in GI microbiota involved in the immune activation. So, when rotenone was administered post-colitis, the intestinal permeability changed more than when rotenone was used alone, making the intestinal milieu more permissive to subsequent inflammatory insult.

Several lines of evidence implicate GFAP^+^ EGC in regulating the gut inflammatory response and the integrity of the gut epithelium^58,59.^ Indeed, our data showed increased enteric glial expression in the myenteric plexuses, supporting the pro-inflammatory state in the colon of mice with disrupted intestinal barrier integrity. In comparison, there was an additive effect due to the administration of rotenone post-colitis, as evidenced by increased enteric glial activation (increased GFAP reactivity in the myenteric plexuses and lamina propria of the colon). These results were further supported by western blot analysis showing the correlation of enteric glia activation with DSS and rotenone. The intestinal microenvironment is intrinsically tied to α-syn production. Several studies have found that rotenone causes *de novo* synthesis of α-syn and can act on primary enteric neurons to secrete α-syn, which is later absorbed and reversely transported to the brain^60^. Our study found that DSS+Rot and Rot alone animals showed higher levels of α-syn expression in their colons, but no effect was observed in DSS alone animals. These events resulted in higher p-syn levels in colonic tissue at 8 weeks of intragastric rotenone post-colitis. These data suggest that, whereas DSS may induce functional damage to the GI, cellular α-syn pathology persists with rotenone exposure.

Epidemiological studies indicate an elevated risk of PD among IBD patients, and gut microbiota plays an essential role in the occurrence of both diseases. Recent evidence also revealed that DSS-induced colitis promotes the development of microbiota dysbiosis^61^. Epidemiological evidence has linked specific pesticide exposures that impact microbiome configuration to varying outcomes^62^. Emerging clinical and animal research using fecal transplantation and germ-free Tg model of PD has now linked imbalances of gut microbiome composition to promote neuroinflammatory response and contribute to enteric α-syn mediated PD progression^63^. At 8 weeks of rotenone administration post-colitis, the microbial community composition of mice from all 3 groups (DSS, Rot, and DSS+Rot) revealed a trend that seemed to separate from control mice. The gut microbiota changes found in colitis and rotenone-treated post-colitis mice in our study are similar to what has been observed in a study on another toxin-based PD mouse model, demonstrating an increasing trend in the RAs of *Bacteroidetes* and a decreased trend in the RAs of *Firmicutes* in PD mice^64–66^. In contrast, other studies show the opposite results^67^. We showed a lower abundance of Lactobacillus at the genus level in DSS alone and DSS+Rot group compared to the control group. This is in line with previous studies that have shown excessive stress caused a shift in microbial composition that resulted in a lower proportion of anti-inflammatory bacteria, such as *Lactobacillus.* These bacteria regulated emotional behavior and reduced disease progression in a mouse model of multiple sclerosis caused by experimental autoimmune encephalomyelitis by inducing transcription of γ-aminobutyric acid receptors^68^. In contrast to the results obtained in our study, Johnson *et al.* reported a significant increase in *Lactobacillus* in the colon of rotenone-treated animals^69^. Variability in microbiota composition between studies may be due to animal species, lack of consistency in rotenone concentration, time, mode of administration, and sample processing and sequencing. Parasutterella genera’s RA showed an increased trend in DSS alone mice, which can be positively correlated with chronic GI inflammation and severity of PD ^70,71.^ In our study, we found a non-significant decrease in RA of *Catenibacterium* in Rot alone and DSS+Rot mice compared to the control group. No studies directly comparing the effects of the abundance of *Catenibacterium* with PD. Still, one recent study reported a reduction of *Catenibacterium*-derived nanovesicles in PD patients, providing a method for diagnosing PD. These vesicles (~100nm in size) are released locally by bacteria, absorbed by the epithelial cells of the mucus membrane, and then transported throughout the body via the lymphatic stream, where they play a role in the immunological and inflammatory response^72^. This might imply that reduced RA of *Catenibacterium* in Rot and DSS+Rot group might be resulted from increased rotenone mediated α-syn toxic species. Notably, a significantly higher level of RA of genera *Turicibactor* were found in new PD patients compared to the corresponding healthy controls^73^. In our study, abundance of *Turicibactor* was dominant at the genus level in DSS+Rot group, while we observed an increased abundance of the genus *Clostridium sensu stricto 1* in the combination group, which has been previously reported to be associated with an elevated risk of PD, identified through genome-wide association studies. Taken together, our findings suggest that remarkable changes in bacterial composition may occur after colitis, whereas rotenone administration after colitis did not affect bacterial species diversity.

Understanding the mechanism underlying neurodegeneration and α-syn accumulation in response to multiple insults is critical. Our *in-vivo* findings indicate that the pre-treatment of DSS induces permeability changes, and microbial dysbiosis positively influences rotenone mediated α-syn deleterious consequences and illustrates the synergy between the action of neuroinflammation and neurodegeneration. Recent evidence in animal models and cell cultures supports the role of α-syn in activating EGC and causing pro-inflammatory cytokines to be released^74^. In our study, rotenone-treated mice showed no signs of colitis compared to the control group, as evidenced by their stable weight throughout the experiment. In contrast to our proposed hypothesis, we found significantly higher TNF-α in the rotenone group compared to the control group, which could be attributed to a rotenone-induced α-syn mediated neuroinflammatory cascade. These results illustrate that rotenone may cause a distinct inflammatory state at long-term exposure^20^. Whereas another active stimulus, DSS, may have provided the necessary intestinal milieu with higher inflammatory cytokines expression (TNF-α, IL-6, IL-1β, and CCL2) for the potential consequences of subsequently administered low-dose rotenone. Importantly, pre-existing DSS-induced inflammation and rotenone-mediated -syn pathology may interact in a feedback loop, allowing further α-syn accumulation and worsening DAergic neuronal death in the colon (as evidenced by a dramatic decrease in TH expression in the DSS+Rot group compared to the control group). However, in the case of DSS and Rot alone, we did not observe DAergic neuronal death in colon relative to control. Combining these finding suggests that distinct intestinal milieu in intestinal nerves of DSS+Rot causes toxic and aggregation-like changes under inflammation, which might be responsible for inducing the death of DAergic neurons.

Numerous shreds of evidence show systemic inflammatory processes exacerbate ongoing neurodegeneration in PD patients and animal models. In the present study, we found that markers of inflammation (TNF-α, IL-6) were not elevated in the blood of DSS+Rot mice at the 18^th^ week, suggesting a non-systemic mode of disease progression. A few findings in rotenone administration in preexisted colitis require additional discussion. First, serum levels of TNF-α were upregulated in mice shortly after colitis induction; however, this upregulation was transient, returning to normal levels during rotenone treatment in later weeks. This possibility is supported by animal studies in which elevation of Serum TNF-α was correlated with colonic injury in the DSS-induced chronic colitis model^75^.

It is becoming increasingly evident that α-syn pathology and degeneration of ChAT^+^ neurons in DMV represent a tipping point in PD progression. Therefore, we examined whether a low dose of rotenone post-colitis-induced neurodegeneration was accompanied by α-syn pathology in DMV. We observed a significant reduction of cholinergic neurons in the DMV of DSS+Rot and Rot alone treated mice compared to control mice. Interestingly, the ChAT^+^ signal in the vagal nerve was unchanged in the DSS alone group, representing functional effect caused by DSS is restricted to the colon only. Moreover, α-syn was accumulated in neuronal soma in the DMV neurons of DSS+Rot and Rot alone mice, is also coincided with observed neurodegeneration. Furthermore, we also observed an inflammatory-associated reaction with increased GFAP in DSS+Rot and Rot alone mice but not control and DSS alone mice. Pan montojo *et. al* previously found that administration of intragastric rotenone for 1.5 months promotes α-syn pathology and loss of ChAT^+^ neurons during PD progression before degenerating DAergic neurons later in the disease. The present data support that idea^20,76.^ Together with these observations, we can say that our rotenone model involves retrograde vagus nerve-mediated propagation of α-syn from GIT to the brain.

It is reported that LC is among the first brain regions affected by α-syn pathology, which promotes LC hyperactivity and non-motor symptoms during PD progression before the degeneration of DAergic neurons later in the disease^77^.: The appearance of α-syn pathology and significant reductions in noradrenaline’s central level in LC is a ubiquitous feature of PD that LC neuronal damage is present as indicated by TH^+^ neuronal cell shrinkage^78,79.^ To determine the effect of intragastric administration of low-dose rotenone post-colitis on LC neurons, we measured the TH and α-syn immunofluorescence intensity. We observed a trend toward increased TH immunoreactivity at the 8^th^ week of intragastric rotenone treatment, but it was not statistically significant which may indicate a compensatory enhancement in TH synthesis. While no significant difference were detected in LC α-syn content at 8^th^ week of rotenone administration. However, in the presence of pre-existed colitis, there was significant reduction in TH intensity, with subsequent increase in α-syn intensity in TH^+^ neurons as compared to control group. Unexpectedly, we also found that trend toward decreased TH in DSS alone treated group. It is likely due to the increased number of GFAP^+^ cells by a possible migration from the adjacent brain areas and bloodstream after the damaging stimulus induced by DSS intoxication, contributing to LC’s regional degeneration in idiopathic PD pathology. Therefore, it is possible that the spread of pathogenic α-syn and endogenous factors could make neurons susceptible to harm in DSS alone animals. Furthermore, our data provide evidence for inflammation occurring in and near the LC of DSS+Rot and DSS alone animals, as confirmed by GFAP^+^ staining around TH^+^ stained neurons in the LC but not outside of the LC. Our findings of inflammation in LC and loss of TH^+^ neurons argue for the loss of nor adrenaline synthesis as contributing cause.

As described, exposure of mice to rotenone causes accumulation of α-syn in enteric and brain stem vagal neurons. This has been involved in the development, and progressive neurodegeneration observed in SNpc. In general, Tg overexpression of human α-syn generates extensive α-synucleinopathies and varying levels of neurodegeneration across many animal species^80^. Only a few α-syn Tg mice models display overt degeneration of nigrostriatal DA fiber, and even fewer indicate loss of nigral DA neurons^81,82.^ The lack of DA neuron loss in most gene-based PD animal models, as well as the presence of α-syn pathology in ENS of most toxin-based PD animal models indirectly by promoting GI inflammation, highlight the importance of enteric α-syn environment interactions in PD. In this line, Martínez *et al.* were the first to check the effect of acute colonic inflammation induced by DSS pre-treatment on MPTP induced Parkinsonism mouse model. Their findings demonstrated that DSS-induced injury circumscribed to the colon exacerbates DAergic neurodegeneration and glial response in the nigrostriatal pathway^83^. Our immunofluorescence experiment results supported previous research and showed co-localization of α-syn in TH^+^ neurons of SNpc in rotenone treated pre colitic mice compared to the control group. DAergic neurons of SNpc appear particularly vulnerable to the effect of α-syn pathology. The above changes are consistent with our finding of reduced TH^+^ neuron count and fibers loss in SNpc and striatum, respectively, in rotenone-treated colitic mice compared to all three groups. Furthermore, we studied glial response under these pathological conditions. We then analyzed astroglia activation by GFAP immunolabeling. In the SNpc and in the striatum, astroglia expression intensity was significantly increased in the DSS+Rot group compared to all three groups. P-syn is a marker of pathological aggregation of α-syn, and we found that expression levels of p-syn in the striatum increased in DSS+Rot group, with concomitant increased gene expression of inflammatory marker (TNF-α, IL-6, IL-8 and CCL2). Overall, the pathological aggregation of α-syn induced by DSS+Rot may play an important role in the gut-brain axis.

These findings are compelling from a translational standpoint since, along with clinical evidence, they clearly show how inflammatory events, even from the gut, and especially if long-lasting, constitute a significant risk for developing or worsening PD presentation. Our findings further reveal that when combined with rotenone-mediated α-syn exposure, PD pathogenesis’s progression behaves entirely differently than chronic inflammation. This highlights the importance of focusing more on managing chronic inflammatory disorders, with the eventual purpose of avoiding α-synucleinopathies and other neurodegenerative diseases.

There are some study limitations worth noting. There is no ideal animal model for PD, and each model recapitulates only some aspects of PD. In this study, we did not clarify the possible pathways by which intragastric rotenone for 8 weeks post-colitis exacerbated the α-syn expression in the colon. Second, the changes in subpopulations of immune cells in the colon are only preliminary studies, and more experiments are needed to demonstrate how the interactions of intestinal and environmental toxins interact with colonic immune cells. Furthermore, DSS was administered in three cycles to mimic chronic, intermittent barrier dysfunction; future research may extend this treatment period to four to five cycles to see if a longer duration is sufficient to induce more severe intestinal barrier dysfunction/inflammation and promote the PD-like phenotype. Another thing to look at is how the response to low-dose DSS may be affected by differences within the group and differences in the microbiota between institutions.

## Declarations

## Acknowledgements

We thank the miBiome Therapeutics LLP (Mumbai, India) for 16S rRNA sequencing service. We also thank RECETOX, Masaryk University (Brno, Czech Republic) for analyzing 16S rRNA sequencing data.

## Conflicts of Interest

The authors declare that there is no conflict of interest.

## Consent for publication

Each author has provided his/her consent for this publication.

## Authorship contribution

We declare that all authors made fundamental contributions to the manuscript. NS, and AMK designed the study. NS, MS, DT, and HK conducted experiments. NS, SS, LB, and PA analyzed the data. NS, and AMK participated in the interpretation and writing of the manuscript. HK and AMK advised the experiments. All authors read and approved the final manuscript.

## Funding

This supplement was supported by the National Institute of Pharmaceutical Education and Research (NIPER) seed fund-Ahmedabad, Ministry of Chemicals and Fertilizers, Government of India. and by project no. LX22NPO5107 (MEYS): financed by European Union – Next Generation EU. Dr. Amit Khairnar gratefully acknowledges the support of the Ramalingaswami Fellowship (No. BT/RLF/Re-entry/24/2017) from the Department of Biotechnology, India.

## Ethical approval

No clinical study was conducted. This study was performed in accordance with CPCSEA guidelines. Approval was granted by IAEC committee of NIPER Ahmedabad (NIPERA/IAEC/2018/032).

## Availability of data and materials

The datasets supporting the conclusions of this article are included within the article and its additional files. All material used in this manuscript will be made available to researchers subject to confidentiality.

## Abbreviations

α-syn: Alpha synuclein
ANOVA: One-way analysis of variance
BCA: Bicinchoninic acid
CMC: Carboxymethylcellulose
CNS: Central nervous systems
ChAT: Choline acetyl transferase
CPCSEA: Committee for the Purpose of Control and Supervision of Experiments on Animals
CCl2: C-C chemokine ligand 2
DAI: Disease Activity Index
DAergic: Dopaminergic
DAB: 3,3′-diaminobenzidine
DAPI: 4′,6-diamidino-2-phenylindole
DSS: Dextran sodium sulphate
DMV: Dorsal motor nucleus of vagus
ENS: Enteric nervous system
EGC: Enteric glial cells
GI: Gastrointestinal
HRP: Horseradish peroxidase
IBD: Inflammatory Bowel Disease
LRRK2: Leucine rich repeat kinase 2
LC-MS: Liquid chromatography mass spectroscopy
LC: Locus coeruleus
OD: Optical Density
p-syn: Phosphorylated S129 synuclein
PD: Parkinson’s disease
PBS: Phosphate buffer saline
PB: Phosphate buffer
PFA: Paraformaldehyde
PMSF: Phenylmethyl sulfonyl fluoride
RA: Relative Abundance
PVDF: Polyvinylidene difluoride
RBW: Round beam walk
SDS: Sodium dodecyl sulfate
SNpc: Substantia nigra pars compacta
Tg: Transgenic
TH: Tyrosine hydroxylase
ZO-1).: Zona Occludens

## Notes

### Competing Interest Statement

The authors have declared no competing interest.

