## Supplementary data for "Intragastric administration of low-dose rotenone post-colitis exacerbates damage to the nigrostriatal dopaminergic system in Parkinson’s disease: The pace accelerates even more"

Author Affiliations:

Co-authors:

Nishant Sharma

Monika Sharma

Disha Thakkar

Hemant Kumar

Sona Smetanova

Lucie Burešová

Petr Andrla

Amit Khairnar

### Supplementary section (S1):

**DNA isolation:** Twelve samples were received from the client labelled as C1, C2, C3, R1, R2, R3, RD1, RD2, RD3, D1, D2,D3 and stored at -80° C. All these samples were given miBiome internal IDs as mentioned in Table 1 below. DNA was isolated using QIAamp PowerFecal Pro DNA Kit, Cat:51804 as per manufacturer's instructions.The DNA samples were quantified on NanoDrop One and Qubit using water and standard as controls respectively. An aliquot of the DNA was electrophoresed on a 0.8% agarose gel in 1X TAE along with DNA ladder, stained with Syber SAFE and visualized in Gel-doc (Biorad).

Based on DNA quantity and integrity criteria, all samples were taken ahead for library preparation for 16S rDNA based metagenomics.

| Client ID on tube | Nanodrop DNA conc. (ng/µl) | A260/280 | A260/230 | Qubit DNA conc (ng/µl) |
|---|---|---|---|---|
| C1 | 210.8 | 1.92 | 0.77 | 132 |
| C2 | 293.2 | 1.89 | 1.31 | 218 |
| C3 | 117.8 | 1.9 | 0.46 | 90 |
| R1 | 127.9 | 1.85 | 1.28 | 105 |
| R2 | 531.3 | 1.91 | 2.17 | 452 |
| R3 | 191.1 | 1.89 | 0.53 | 154 |
| RD1 | 682.6 | 1.88 | 2.22 | 568 |
| RD2 | 399 | 1.91 | 2.11 | 330 |
| RD3 | 419.8 | 1.91 | 1.44 | 316 |
| D1 | 532.8 | 1.91 | 2.25 | 388 |
| D2 | 352.5 | 1.9 | 0.54 | 308 |
| D3 | 497.1 | 1.92 | 2.1 | 400 |

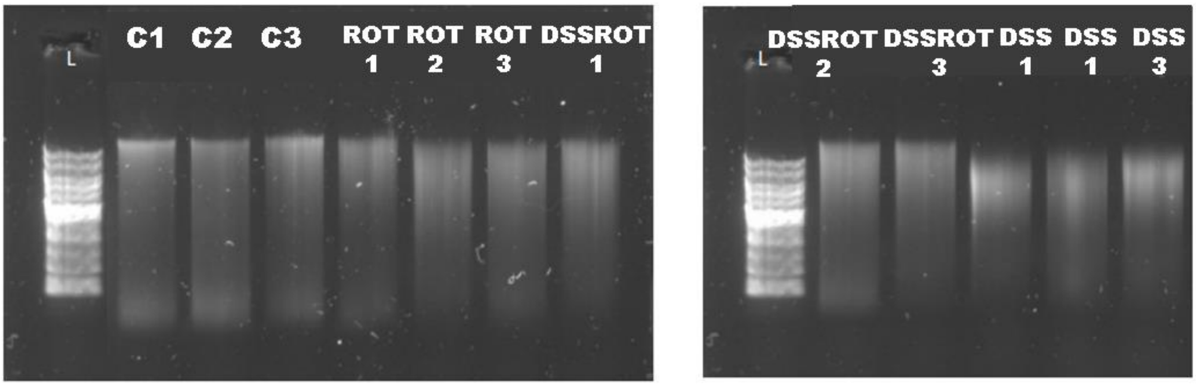


**Figure1**: Agarose gel profile of DNA extracted from mice fecal samples co-electrophoresed with 1 kb DNA ladder (lane L).

***16S rRNA gene sequencing bioinformatical analysis:***

Paired reads were first processed using an in-house pipeline implemented in Python 3. Steps of processing included the trimming of low-quality 3' ends of reads, removal of read pairs containing unspecified base N, and removal of pairs containing very short reads. In order to minimize sequencing and PCR-derived error, forward and reverse reads were denoised using the DADA2 amplicon denoising R package[1]. Following denoising, the forward and reverse reads were joined into a single longer read using the fastq-join read joining utility [2]. In order to be joined, reads in pairs had to have an overlap of at least 20 base pairs with no mismatches allowed. As the final step, chimeric sequences were removed from the joined reads using the remove Bimera function of the DADA2 R package. Subsequent taxonomic assignment was conducted by the uclust-consensus method from the QIIME [3] analysis framework using the Silva v. 132 reference database[4].

**Supplementary section (S2):**

**LC–MS Method:**

The LC-MS study was carried out using an Agilent 1290 ultra-high-performance liquid chromatography system (UHPLC; Agilent technologies, USA) linked to an Agilent 6545 series electro spray ionization-quadrupole-time of flight mass analyzer (ESI-Q-TOF; Agilent technologies, USA). Initially, LC-MS settings were optimised for rotenone in positive mode scan. The chromatographic separation was accomplished on a Poroshell 120 SB C-18 (3100 mm, 1.8 m, Agilent Infnity Lab, USA) column with a mobile phase of methanol (solvent A) and 0.1% formic acid (solvent B). The gradient programme was set as follows: 0 min, 70% B; 1 min, 70% B; 3 min, 30% B; 5.5 min, 5% B; 7.9 min, 5% B; 9 min, 70% B; and 10 min, 70% B. The autosampler temperature, flow rate, and injection volume were all set to 15 °C, 0.5 mL min1, and 5 L, respectively. The analyte's MS conditions were optimised in tuning mode. The optimum ESI source settings were as follows: 160 V fragmentor volt age, 3000 V capillary voltage, 65 V skimmer, and 1000 V nozzle voltage. Nitrogen was employed as a drying gas at a flow rate of 10 L min1 at 350 °C.

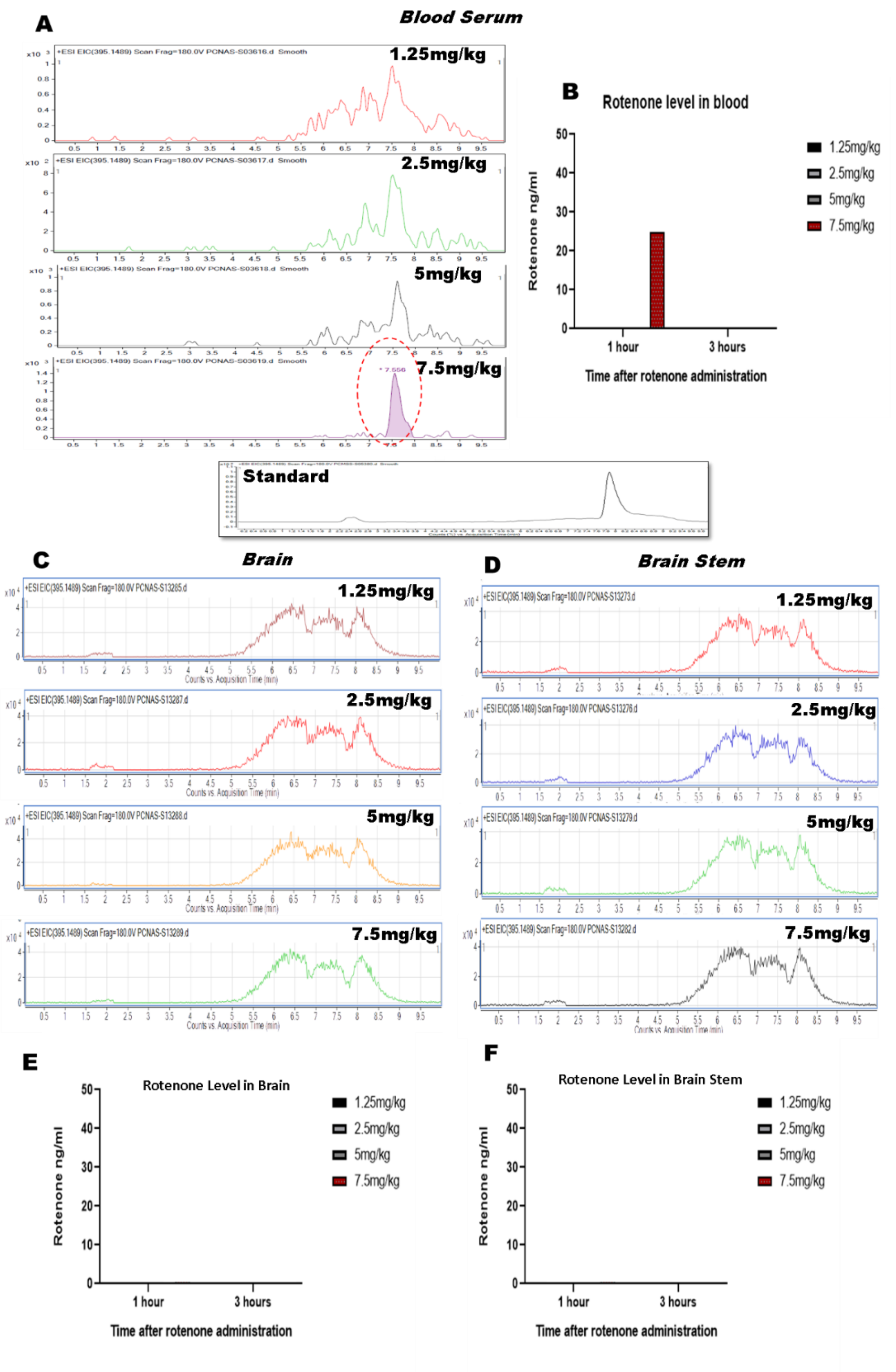


**Figure 1:** A, C and D. Standard (50 ng/ml) and chromatogram from blood serum, Brain and brain stem samples of 1.25, 2.5, 5 and 7.5 mg/kg intragastric rotenone treated for one week after the induction of coitis in mice. B, E and F. Quantification of rotenone levels in blood serum, Brain and brain stem samples collected at 1 and 3 hours after treatment with different doses on 7$^{th}$ day of treatment.

**<u>Supplementary sections 3</u>:**

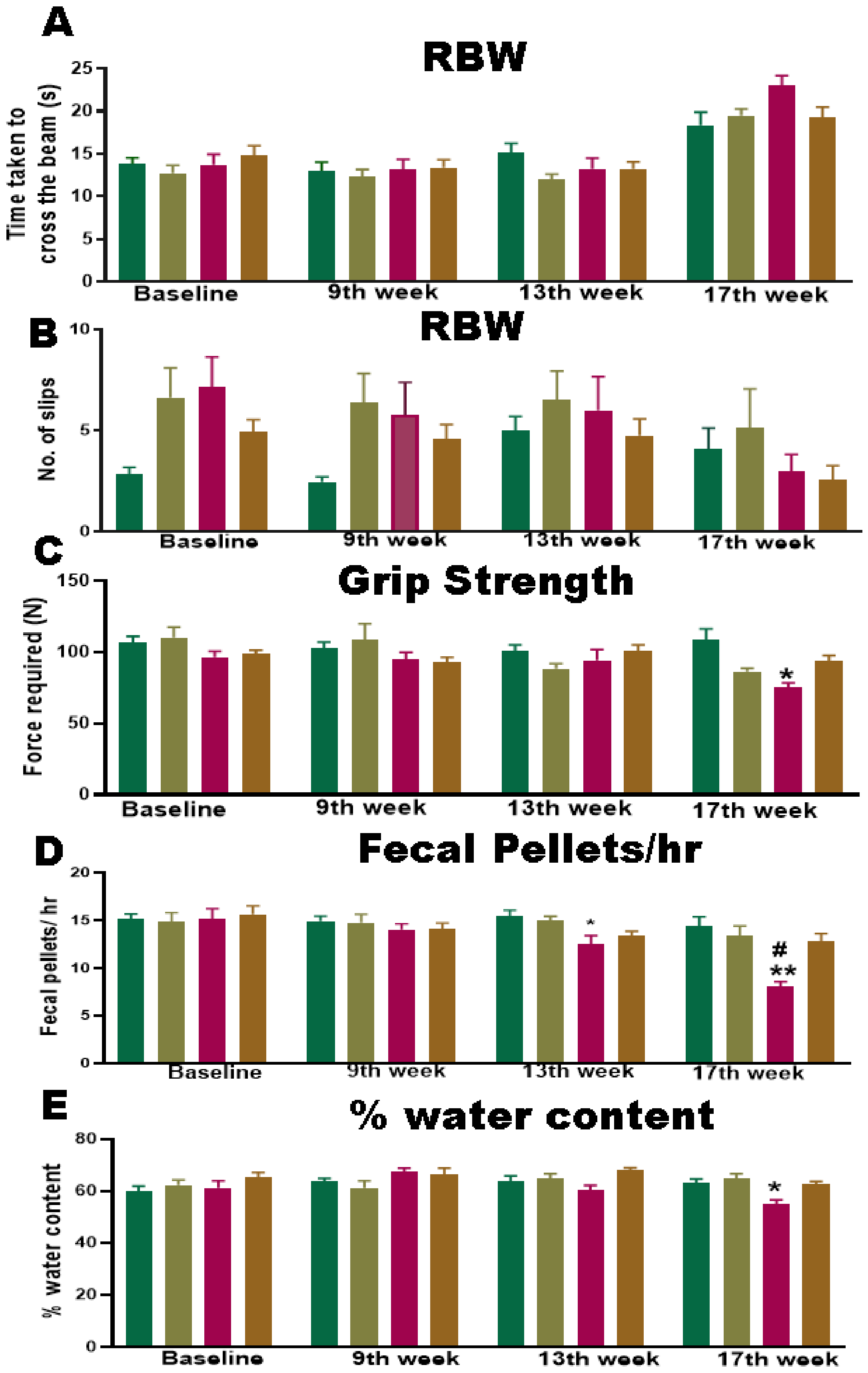

**Figure 3:** Effect of low dose rotenone starting from Baseline, 9th, 13th, and 17th on behavioral and GI parameters in post colitis mice model. A) Time taken to cross the beam, B) Number of foot slips are from the round beam walk test C) Muscle strength in grip strength test, D) No. of fecal pellets per hour, and E) Percent water content in fecal samples. Data is presented as Mean ± SEM (n=5-6). *p < 0.05 vs control group, #p < 0.01vs rotenone group. Statistical analysis was performed using one-way analysis of variance (ANOVA) followed by Tukey's test.

### Supplementary section (S4):

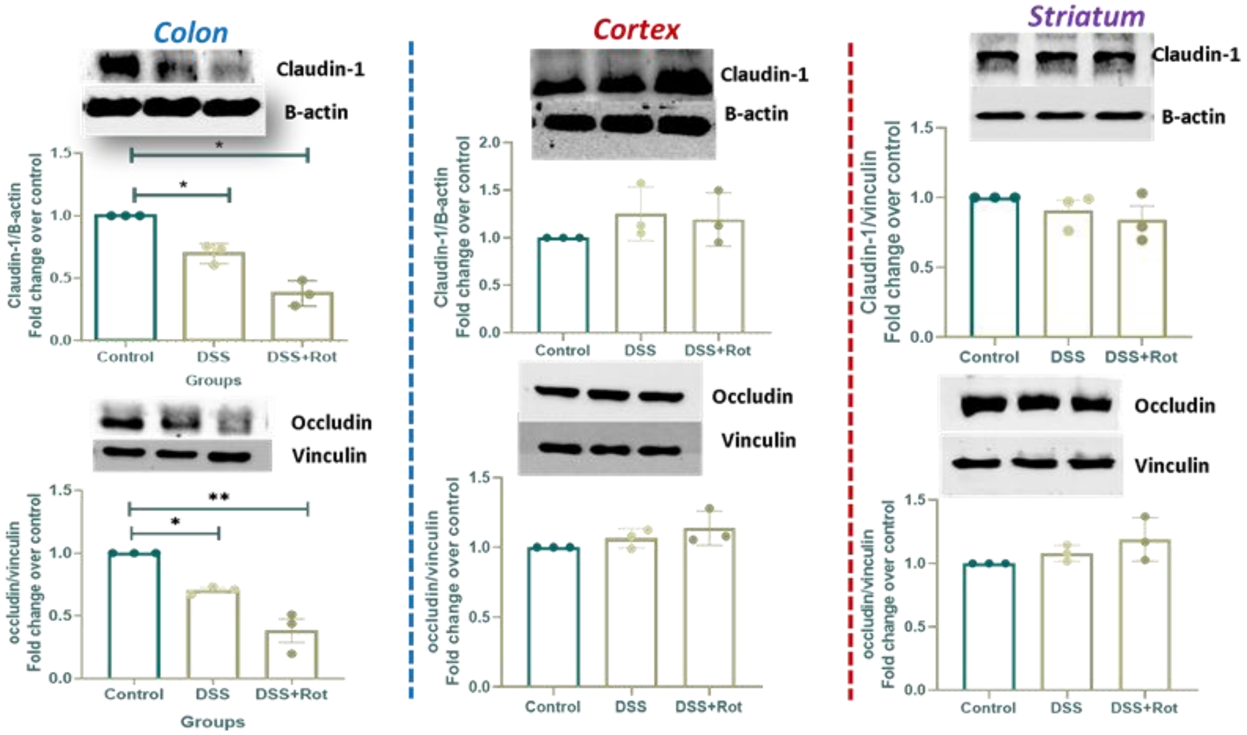


**Figure 4:** Representative western blot image and quantification graph showing the effect of low dose administration of rotenone on protein expression of, Claudin-1 and occludin in colon, striatum and cortex of chronic colitis mouse model as found in pilot study. The results were expressed as Mean ± SEM (n = 3). *p<0.05, ** p < 0.01, vs control group. Statistical analysis was performed using one-way analysis of variance (ANOVA) followed by Tukey's test

1. Callahan BJ, McMurdie PJ, Rosen MJ, Han AW, Johnson AJA, Holmes SP. DADA2: High-resolution sample inference from Illumina amplicon data. *Nature methods*. 2016;13(7):581-583.
2. Aronesty E. Comparison of sequencing utility programs. *The open bioinformatics journal*. 2013;7(1)
3. Caporaso JG, Kuczynski J, Stombaugh J, et al. QIIME allows analysis of high-throughput community sequencing data. *Nature methods*. 2010;7(5):335-336.
4. Quast C, Pruesse E, Yilmaz P, et al. The SILVA ribosomal RNA gene database project: improved data processing and web-based tools. *Nucleic acids research*. 2012;41(D1):D590-D596.
